# Microbial inoculation accelerates post-fire soil recovery in a mixed conifer forest

**DOI:** 10.64898/2026.06.12.731757

**Authors:** Elliot L. Weiss, Jillian F. Banfield

## Abstract

High-severity wildfires of increasing size and frequency result in release of carbon dioxide and loss of timber resources, reduction in biodiversity, loss of soil, diminished water quality, and reduced recreational opportunities. Forest recovery strongly depends on the reestablishment of soil microbial communities, motivating research on how restoration of soil microbiomes in burned forests can be accelerated. Here, we used a high intensity burn pile experiment to test the effectiveness of post-fire native soil amendment. This design enabled us to sequentially and simultaneously sample unburned, burned, and inoculated burned soils while holding post-fire abiotic factors constant. All conditions were sampled at six time points across an annual hydrological cycle and analyzed using 16S and ITS rRNA amplicon sequencing, genome resolved metagenomics, metatranscriptomics, and soil chemistry. Fire sharply reduced bacterial and fungal diversity and eliminated ectomycorrhizal and ericoid symbiotic fungi. Inoculating the burned soil with native microbes accelerated recovery of microbial diversity and of functions associated with nutrient cycling, especially nitrogen transformations. Despite introducing the full diversity of soil microbes from adjacent unburned forest, only a small subset of adapted organisms were engrafted. Native soil inoculation stimulated reestablishment of mycorrhizal fungi, including genera that form essential symbioses with conifers, although this response was not persistent over the full year. Nonetheless, reestablishment of mycorrhizal fungi for even a window of time may facilitate early forest regrowth. We conclude that, by microbial inoculation, recovery that would otherwise rely on dispersal from distant sites is accelerated, potentially enhancing reforestation efforts.

## Introduction

High-severity wildfires are among the most ecologically disruptive forces shaping terrestrial ecosystems. These effects are compounding: the climate changes that are driving more frequent and severe wildfires^1,2^ can also slow or limit post-fire recovery, raising the potential for self-reinforcing cycles of ecosystem degradation and regime shift in fire-prone landscapes^3,4^. Severe burns, which can reach or exceed 500°C at the soil surface^5^, also typically alter soil carbon stocks and aggregate structure, increase surface hydrophobicity^5^, and cause substantial losses of soil nitrogen through volatilization^6^, collectively heightening erosion risk, reducing water infiltration, and increasing the risk of converting productive forest to non-forest land^5^. Beyond the immediate loss of above-ground biomass, high temperatures during intense burns partially sterilize the upper soil horizon, greatly reducing living microbial biomass^7^ and diversity^8–10^. This is important because the soil microbiome plays a central role in the biogeochemical processes that sustain forest productivity. Bacteria, fungi, and archaea mediate nutrient cycling and decomposition^11^, including the breakdown of recalcitrant organic matter and release of plant-available nutrients^12^, and produce plant-stimulating phytohormones^13^.

Ectomycorrhizal fungi (EMF) form essential symbioses with most temperate and boreal forest trees, extending root access to nutrients and water while channeling photosynthate-derived carbon into mineral soil, enhancing stress and pathogen resistance, and promoting plant growth^14^. In fact, it is estimated that around 90% of plants have beneficial root-associated fungi^15^.

Recovery of the soil microbiome to pre-disturbance community composition following high-severity wildfire is a slow process. Studies tracking burned soils over multi-year to multi-decadal periods suggest that bacterial diversity can partially rebound on the order of years, yet fungal communities, particularly EMF, may require many decades to approach compositional equivalence with unburned soils^16–18^. This trajectory is largely driven by succession: early post-fire soils are dominated by fire-loving (pyrophilous) and other pioneering (ruderal) taxa that differ markedly from the mutualistic communities associated with mature ecosystems^19^. Natural microbiome reassembly following high-severity fires depends on stochastic inputs of microbial propagules from surrounding unburned areas via wind, hydrological events, and migration from deeper soils^20^. The effectiveness of wind and water dispersal, particularly for fungi, will decrease as distance from unburned soil increases, so immigration across large burn patches is likely to be very slow^21–23^.

Ecological restoration science has explored the use of soil inocula to accelerate microbiome reassembly in degraded habitats^24–26^, with promising results. Studies pertaining to broadcasting whole soil for bioremediation are less common, yet, the principle is straightforward: by introducing a diverse microbial propagule bank, inoculation bypasses the stochastic dispersal-limited early stages of succession. However, although organic and mineral amendments (e.g., mulch, biochar, compost) have been applied in burned forests^27^, the use of whole-soil inocula to steer soil microbiome recovery in severely burned conifer forests remains untested. To our knowledge, no controlled field study has yet evaluated whether whole-soil inoculation can measurably redirect microbiome trajectories in severely burned conifer forest soils.

Here we report the results of a 12-month, multi-omic field experiment designed to test whether inoculation with adjacent unburned forest soil can accelerate microbiome recovery in high-severity burned plots. To emulate realistic high-severity fire while maintaining environmental control, we constructed large burn piles from locally sourced mixed-conifer logs within the canopy at Blodgett Forest Research Station (El Dorado National Forest, California) and burned them to peak surface temperatures emulating the intensity of high-severity wildfires^28^. Burned subplots were assigned to inoculation with pooled, unburned soil (n = 3) or to untreated burned controls (n = 3), and were sampled across six time points (T0–T5) spanning one year. We integrated 16S rRNA amplicon sequencing (bacterial communities), ITS amplicon sequencing (fungal communities), shotgun metagenomics (metabolic potential and metagenome-assembled genomes, MAGs), and metatranscriptomics (active community gene expression) to characterize (i) trajectory of community composition and diversity in each condition, (ii) genomic determinants of community assembly and succession, (iii) the persistence and activity of introduced microbial taxa, and (iv) the functional consequences of inoculation for soil carbon and nitrogen cycling. Our results show that inoculation transiently elevates bacterial and fungal diversity, promotes the establishment of EMF for an extended period, and shifts functional gene expression toward pathways associated with organic matter decomposition and nitrogen mobilization. Thus, native soil inoculation can accelerate convergence toward an environmentally determined recovery endpoint, with clear relevance to post-fire restoration practice.

## Results

### Site conditions, burn validation, and soil chemistry

A high-intensity experimental pile burn was constructed and burned within the canopy of the mixed conifer forest at Blodgett Forest Research Station (Fig. S1A). Peak soil temperatures reached 800°C at the soil surface during the burn and exceeded 200°C for approximately 4 days, consistent with high-severity wildfire conditions. Following extinguishment, the burn footprint was subdivided into 1 × 1 m subplots assigned to inoculated or untreated burn conditions (Fig. S1B); adjacent intact forest plots served as unburned controls. Sampling spanned November 2023 through December 2024, capturing six time points (T0–T5; T0 equating to time of inoculation, approximately 2 weeks post-fire) across the full annual hydrological cycle (Fig. S2). This period encompassed early and late winters and a dry summer, providing a climatically realistic context for evaluating microbiome recovery and inoculation effects. Volumetric soil moisture closely tracked precipitation, with unburned soils retaining higher moisture at both 0–5 cm and 5–10 cm depths than burned or inoculated plots across most time points; this difference diminished during the late-summer dry period at T4, when soil moisture converged across conditions (Figs. S3A and S3B).

At T0, the burned soil profile showed a visible dark, ash-laden surface horizon overlying lighter mineral soil (Fig. S1C), and soil pH at 0–5 cm was significantly elevated in burned plots relative to unburned soils (7.16 ± 0.28 vs. 6.03 ± 0.36; two-sample t-test, t = 4.28, p = 0.013; Fig. S3C), indicating substantial ash inputs. Inoculated plots also exhibited elevated pH at T0 (6.70 ± 0.12), consistent with shared ash inputs across the burn footprint. In contrast, pH at 5–10 cm did not differ significantly between burned and unburned soils at T0 (6.57 ± 0.36 vs. 6.16 ± 0.40; two-sample t-test, t = 1.31, p = 0.26; Fig. S3D). pH in burned and inoculated plots declined steadily over the 12-month period, nearly converging with unburned values by T5, consistent with progressive leaching and biotic recovery of acidifying processes.

Soil organic carbon and total nitrogen did not differ significantly between burned and unburned soils at T0 (p > 0.05), partly due to high spatial heterogeneity in the unburned plots. In contrast, nitrogen and ash-associated ions showed strong fire effects. At 0–5 cm, NH₄⁺ concentrations were substantially higher in burned than unburned soils (3.86 ± 1.90 vs. 0.07 ± 0.13 ppm; two-sample t-test, t = 3.44, p = 0.026), indicating a pronounced ammonium pulse at the surface. NH₄⁺ was significantly elevated at 5–10 cm in burned soils relative to unburned as well (3.87 ± 1.26 vs. 0.00 ± 0.00 ppm; two-sample t-test, t = 5.41, p = 0.006), demonstrating that the ammonium pulse extended into the subsurface. Manganese was significantly elevated in burned compared to unburned plots at T0 at both 0–5 cm (204.13 ± 51.06 vs. 43.03 ± 43.35 ppm; two-sample t-test, t = 4.17, p = 0.014) and 5–10 cm (195.77 ± 50.24 vs. 39.47 ± 50.61 ppm; t = 3.80, p = 0.019), consistent with ash-derived metal inputs into the soil profile. The results confirm that experimental plots entered the study in a chemically altered state characteristic of severe fire disturbance.

### Microbial community composition

At T0, the time of inoculation, burned soils were compositionally dominated by bacteria from a small number of phyla, principally Pseudomonadota, Actinomycetota, and Bacillota, characteristic of the early post-fire successional stage of copiotrophic and stress-tolerant generalists (Fig. 1). In contrast, unburned soils exhibited greater phylum-level diversity throughout the study. The inoculum that was added to some burned soil plots was a homogenized mixture of unburned soil sampled from adjacent to the burn piles and unburned plots and compositionally very similar to unburned soils in control plots (Fig. S4). Inoculated plots showed greater phylum-level diversity relative to burn-only plots from early timepoints, with partial recovery of Acidobacteriota, Verrucomicrobiota, Planctomycetota, Gemmatimonadota, Myxococcota, and Chloroflexota towards levels observed in unburned soils over the sampling period (Fig. S5).

**Figure 1.**
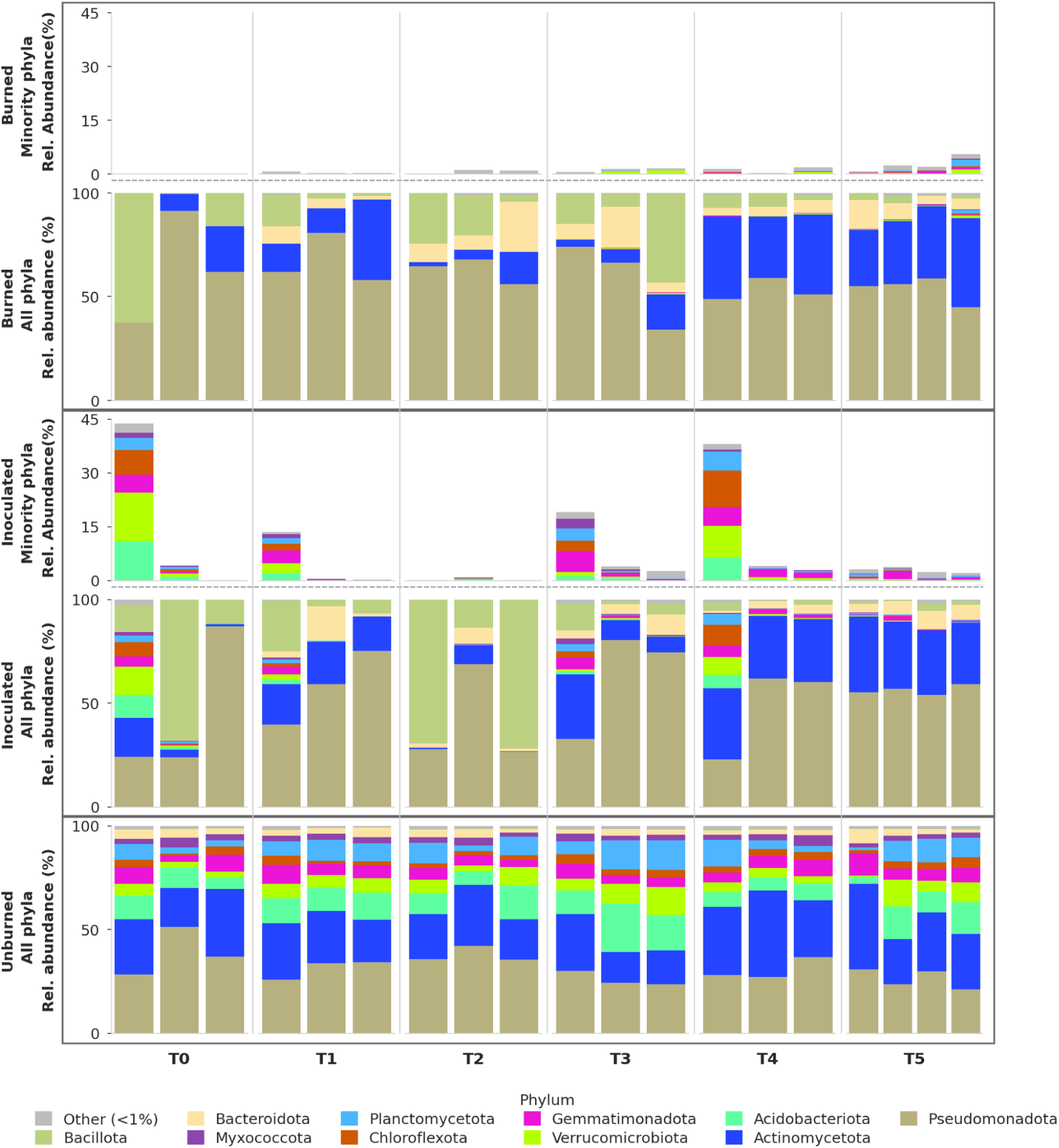
Bacterial community composition at the phylum level across treatments and time points at 0–5 cm depth, based on 16S rRNA amplicon sequencing. Each bar represents one subplot replicate (n = 3 per treatment at T0–T4, n = 4 at T5). For post-burn conditions, the lower panel of each box shows relative abundance (0–100%) of all phyla, and the upper panel shows the same samples with dominant post-burn phyla (Pseudomonadota, Actinomycetota, Bacteroidota, Bacillota) excluded to highlight minority phyla at an expanded scale (0–45%). Burned plots were dominated by Pseudomonadota, Actinomycetota and Bacillota throughout the study period, with Bacteroidota becoming abundant after T0, with limited representation of oligotrophic phyla characteristic of mature forest soils. Inoculated plots showed substantially higher relative abundance of minority phyla from T0 onward, with Acidobacteriota, Verrucomicrobiota, Planctomycetota, and Chloroflexota recovering toward levels observed in unburned soils over the sampling period. Unburned soils maintained a compositionally diverse community throughout.

Bacterial and fungal communities were both compositionally distinct among all conditions at the soil surface (bacterial PERMANOVA condition R² = 0.378, full model R² = 0.65, p < 0.001; fungal condition R² = 0.210, full model R² = 0.45, p < 0.001; n = 57 per kingdom; Fig. S6). A significant condition × timepoint interaction for surface bacterial communities (R² = 0.135, p = 0.02) indicated divergent condition trajectories over time. Condition effects were also significant at depth (5–10 cm) but with reduced effect sizes (bacterial R² = 0.264, p < 0.001; fungal R² = 0.154, p < 0.001), consistent with fire impact attenuation with soil depth. Burned and inoculated plots showed significantly greater among-sample heterogeneity than unburned soils at both depths and for both kingdoms (PERMDISP all p < 0.001), while heterogeneity did not differ significantly between burned and inoculated plots.

Fire significantly and persistently reduced bacterial and fungal alpha diversity at both depths, with inoculated plots trending intermediate but statistically indistinguishable from burned plots (Fig. S7). An unexpected enrichment of Chloroflexota, Acidobacteriota, and Planctomycetota occurred at 5–10 cm in uninoculated burned plots at T4 (Fig. S8). Given the limited sensitivity of community-wide diversity indices to ecologically meaningful compositional change in heterogeneous forest soils^29^, subsequent analyses focus on compositional, functional, and genome-resolved metrics.

### Fire drives a mycorrhizal-to-saprotrophic shift and inoculation restores mycorrhizal communities

Using FUNGuild, we grouped ITS amplicon sequences based on predicted general fungal lifestyles (e.g., ectomycorrhizal fungi (EMF), saprotrophs) and revealed a dramatic restructuring of the fungal lifestyle composition following fire (Fig. 2A). Unburned soils were characterized by a diverse assemblage in which EMF and ericoid mycorrhizal fungi (ErM) jointly dominated, reflecting the intact symbiotic network supporting the mixed conifer overstory and ericaceous (acid adapted) plant understory.

**Figure 2.**
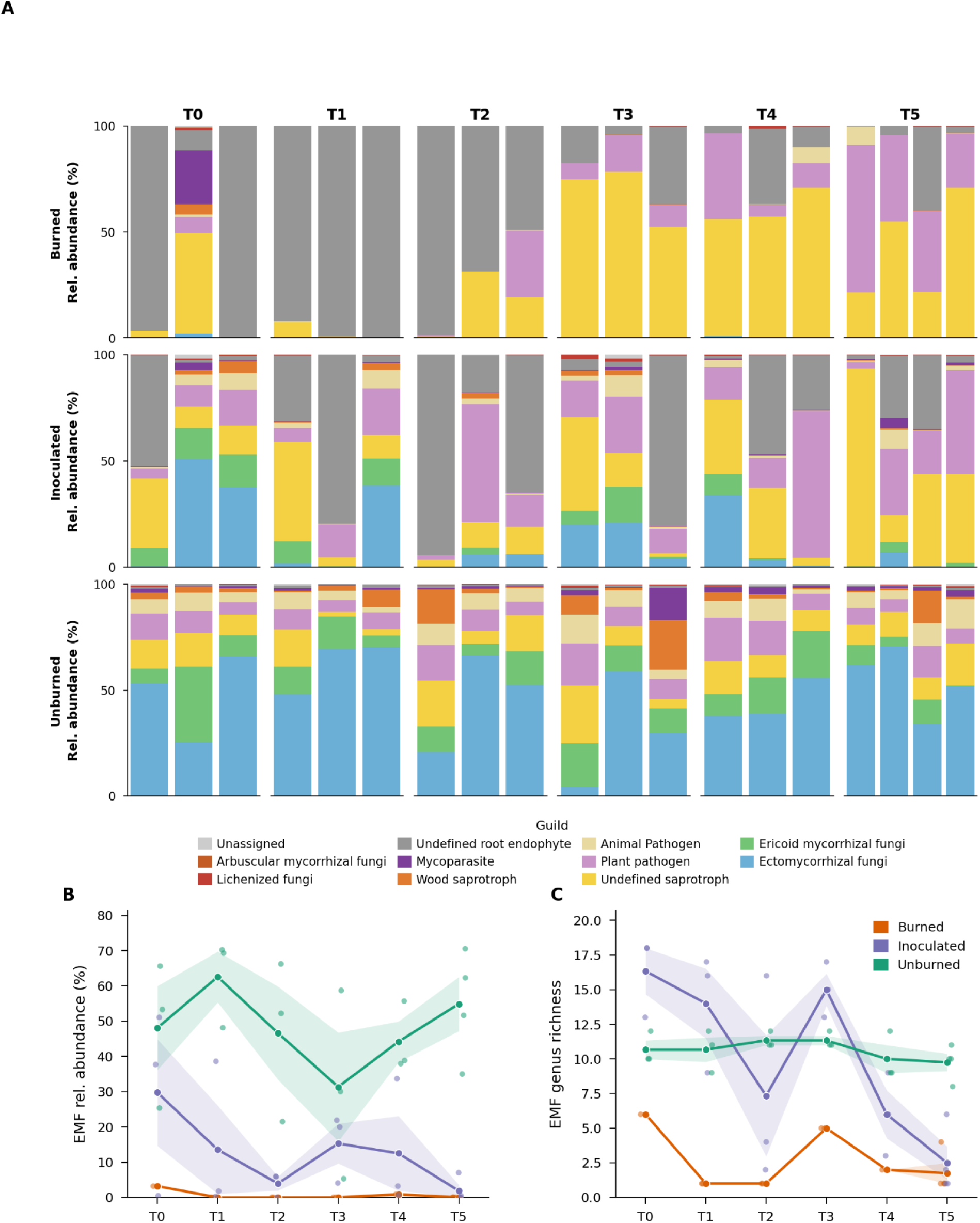
Fungal guild composition and ectomycorrhizal community dynamics across treatments at 0–5 cm depth. (A) Stacked bar plots show relative abundance for each subplot replicate (n = 3 per treatment at T0–T4, n = 4 at T5), estimated from ITS amplicon sequencing with FUNGuild annotation (Probable and Highly Probable assignments only). Unburned soils are dominated by ectomycorrhizal (EMF, blue) and ericoid mycorrhizal (ErM, green) fungi throughout the sampling period. Burned plots are devoid of mycorrhizal fungi at all time points, dominated instead by undefined saprotrophic and root endophyte taxa. Inoculated plots show restoration of EMF and ErM guilds from T0, with abundance variable across the sampling period. (B) Mean EMF relative abundance (as a percentage of all fungal reads) across treatments over the 12-month study period. Inoculated plots maintained EMF relative abundance of 2–30% throughout, compared to near-zero values in burned plots. Unburned soils maintained EMF at 31–63% across all time points. (C) Mean EMF genus richness per subplot across treatments and time points. Inoculated plots supported 6–16 EMF genera throughout the study period, comparable in diversity to unburned soils (10–11 genera) and substantially exceeding burned plots (1–6 genera). Lines connect treatment means; shaded bands indicate ±1 standard error; individual points show per-subplot values.

Burned plots at T0 were almost entirely devoid of mycorrhizal fungi at both depths (Fig. 2A and S9), with the post-fire fungal community dominated instead by saprotrophic taxa capable of colonizing carbon-rich pyrogenic substrates. Burned plots also featured animal and plant pathogenic fungi, with plant pathogens increasing in relative abundance over time. Root endophytes persisted in burned plots in early post-fire samples but declined progressively across the sampling period.

In striking contrast, the post-burn inoculated plots showed a clear return of both EMF and ErM fungi from T0 onward at comparable relative abundance at both 0–5 and 5–10 cm depths. Both EMF and ErM relative abundance was dynamic across the sampling period and variable among subplots, with a notably lower signal at T5. However, EMF relative abundance and genus richness were substantially elevated in inoculated compared to burned plots until the final timepoint (Figs. 2B and 2C), with dominant genera including *Russula*, *Tomentella*, *Rhizopogon*, *Inocybe*, *Helvella*, and *Byssocorticium* (Fig. S10), mirroring the EMF assemblage of the surrounding unburned forest.

### Genome-resolved evidence for persistent r-selection in burned soils

Metagenomic assembly and binning across all samples yielded 1,952 MAGs after dereplication at ≥ 99% ANI, of which 1,502 were draft quality and 450 were near complete (Table S1). After dereplication at ≥ 95% ANI, there were 1,123 MAGs, of which 753 were draft quality and 370 were near complete. The genomes are from 19 bacterial phyla and one archaeal phylum. The species-level (95% ANI) clusters were used for community-level growth-rate and life-history analyses, while strain-level (99% ANI) resolution was used for MAG-resolved functional and biogeochemical cycling analyses and metatranscriptome mapping.

To characterize shifts in the selective regime imposed by fire and inoculation beyond compositional change, we estimated genome-resolved maximum growth rates using codon usage bias for all recovered MAGs^30^ and examined the relationship between estimated doubling time and relative abundance across conditions, time points, and soil depths (Figs. 3A, 3B and S11).

**Figure 3.**
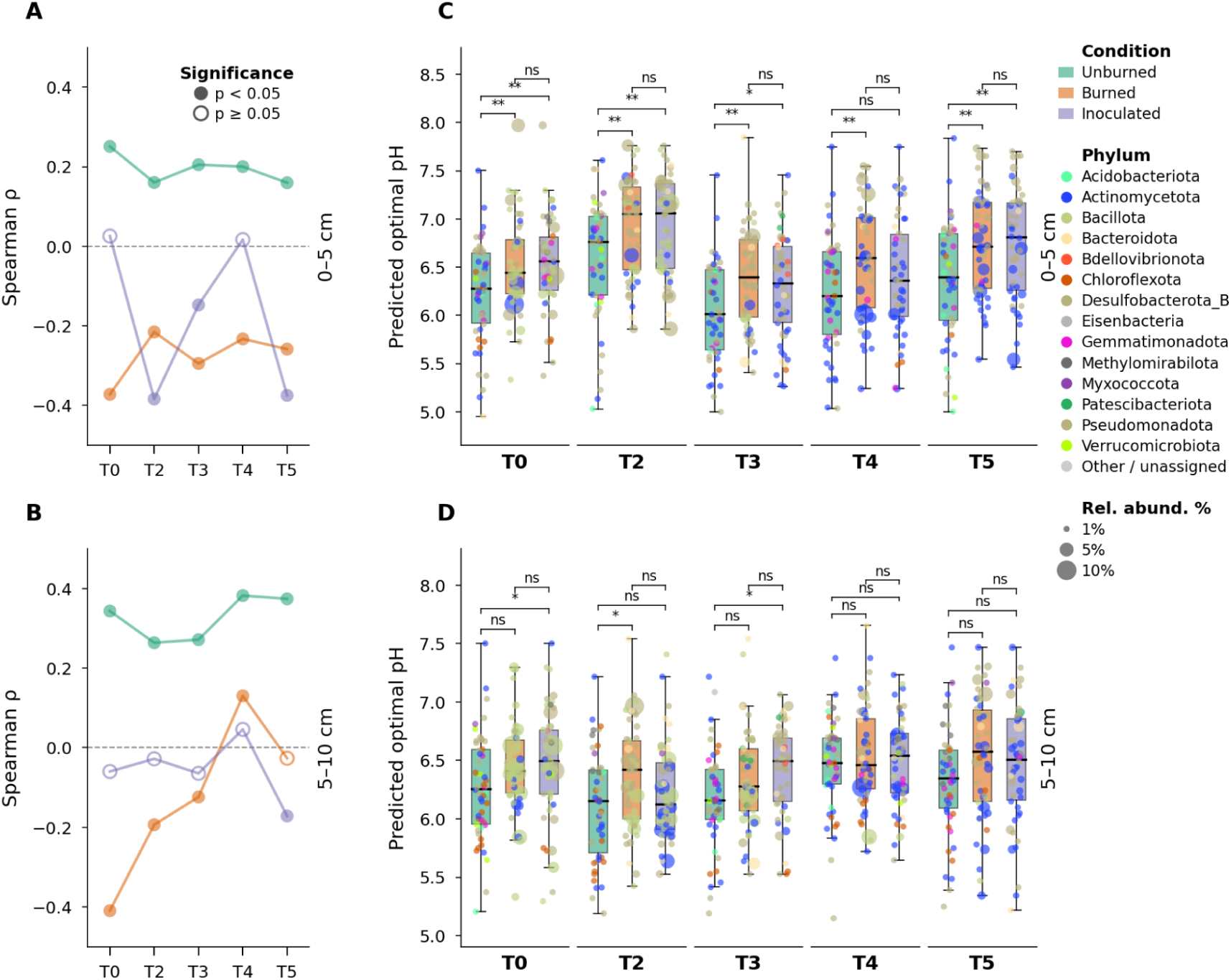
Genome-resolved microbial life-history strategy: growth rate–abundance relationships and predicted pH optima across treatments and depths. (A, B) Spearman correlation coefficient (ρ) between estimated MAG (95% ANI) doubling time and relative abundance at 0–5 cm (A) and 5–10 cm (B), plotted over time for burned, inoculated, and unburned treatments. Positive ρ indicates K-selection (slow-growing specialists dominant); negative ρ indicates r-selection (fast-growing generalists dominant). Filled points indicate p < 0.05; open points indicate p ≥ 0.05. Unburned soils maintain consistently positive correlations throughout the study at both depths (ρ = 0.16–0.38 surface; ρ = 0.25–0.39 subsurface). Fire imposes a persistent shift toward negative ρ at the surface (ρ = −0.22 to −0.37) for the full 12-month period, while subsurface burned communities show a more dynamic recovery trajectory. Inoculated surface plots oscillated between significant r-selection (ρ = −0.18 to −0.4 at T2, T3, T5) and non-significant neutral values (T0 and T4), indicative of an intermediate, dynamic selective regime distinct from the persistent r-selection of burned plots. (C, D) Predicted optimal growth pH derived from the 50 most abundant MAGs (95% ANI) per timepoint at 0–5 cm (C) and 5–10 cm (D), estimated using a machine learning pH preference model. Points are colored by phylum and scaled by relative abundance. Burned and inoculated soils exhibit significantly higher predicted pH optima than unburned soils at the surface across most timepoints (excluding T4, Tukey HSD, p < 0.05), consistent with persistent alkaline selection pressure in post-fire soils. This pattern is attenuated at 5–10 cm depth, where fewer significant differences were detected.

Positive Spearman correlations imply that slower growing organisms dominate, while negative correlations suggest the environment is selecting for organisms capable of faster growth. In unburned soils, a consistent positive Spearman correlation between relative abundance and doubling time at 0–5 cm (ρ = 0.16–0.25, p < 0.001) and at 5–10 cm (ρ = 0.26–0.38, p < 0.001) indicated that slow-growing taxa achieve competitive dominance – a signature of K-selection in stable, resource-limited environments^31^. Burned plots at the soil surface exhibited strongly negative correlations between microbial relative abundance and doubling time at T0 (ρ = −0.37, p < 0.001), demonstrating that fast-growing opportunists dominated the immediate post-fire community. This effect was persistent, as burned surface soils maintained negative correlations throughout the full 12-month study (ρ = −0.22 to −0.37, p < 0.001). Subsurface burned communities (5–10 cm) showed substantially faster return to dominance by K-strategists, with Spearman ρ returning from −0.41 at T0 (p < 0.001) to near zero by T5 (ρ = −0.03, p = 0.51), consistent with more buffered abiotic conditions at depth (Fig. 3B). Inoculated surface plots oscillated between significant r-selection (ρ = −0.18 to −0.4 at T2, T3, T5) and non-significant neutral values (T0 and T4; Fig. 3A), indicative of an intermediate, dynamic selective regime distinct from the persistent r-selection of burned plots.

### Microbial pH growth optima reflect persistent post-fire soil chemistry

Complementing the growth rate analysis, optimal pH for growth was estimated from all MAGs and plotted for the 50 most abundant organisms per timepoint and condition. Dominant bacteria in 0–5 cm burned soils were predicted to have significantly higher optimal growth pH than bacteria in unburned soils at most timepoints (excluding T4; Fig. 3C; p < 0.05; Tukey-adjusted). Inoculated soils also harbored bacteria with significantly higher predicted optimal growth pH than unburned soils at all timepoints except T4, where inoculated soils were intermediate and not statistically distinguishable from either burned or unburned soils (Fig. 3C; p < 0.05; Tukey-adjusted). At 5–10 cm the pattern was attenuated (Fig. 3D). Elevated optimal pH for growth is consistent with the observed increase in soil pH following fire, and the attenuation at depth with reduced direct ash deposition in the subsurface horizon.

### Evidence for accelerated nitrification in inoculated plots

Unburned soils maintained near-zero NH₄⁺ concentrations throughout the study period, reflecting steady-state nitrogen cycling in an intact soil community. NH₄⁺ concentrations were substantially elevated in burned plots relative to unburned plots from T0 onward (Fig. 4A). A notable divergence between burned and inoculated conditions emerged at T3, when inoculated plots showed a marked reduction in NH₄⁺ while burned plots remained elevated (0.71 ± 0.27 vs. 2.31 ± 0.74, respectively, two-sample t-test, p = 0.025). Subsequently, soil NO₃⁻ concentrations exhibited a spike in inoculated plots (at T4), consistent with renewed biological nitrification converting NH₄⁺ to NO₃⁻. Burned plots showed a modest and variable increase in NO₃⁻ concentrations at T4 (Fig. 4B).

**Figure 4.**
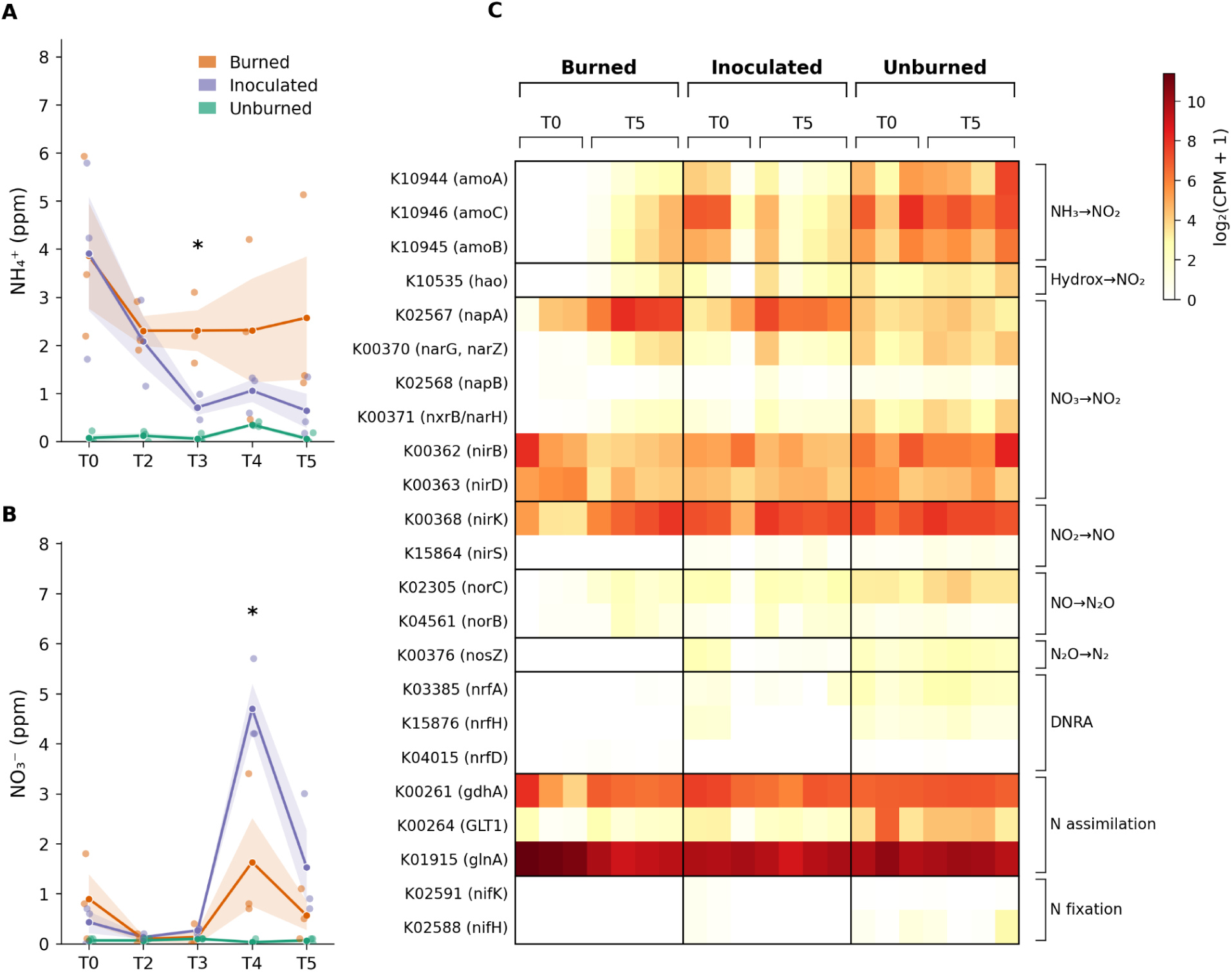
Soil inorganic nitrogen dynamics and metatranscriptomic nitrogen cycling gene expression across treatments at 0–5 cm depth. (A) Soil ammonium (NH₄⁺, ppm) and (B) soil nitrate (NO₃⁻, ppm) measured at T0 and T2–T5 for burned, inoculated, and unburned plots. Lines connect treatment means; shaded bands indicate ±1 standard error; individual points show per-subplot values (n = 3 per treatment). Stars above timepoints indicate significant differences between burned and inoculated plots only (two-sample t-test assuming equal variances, two-sided; unburned significance levels not shown for clarity). NH₄⁺ concentrations were substantially elevated in burned and inoculated plots relative to unburned soils from T0 onward. Inoculated plots showed a decline in NH₄⁺ by T3 (April 2024), diverging from burned plots which remained elevated throughout the study period (0.71 ± 0.27 vs. 2.31 ± 0.74, respectively, two-sample t-test, p = 0.025). A pronounced NO₃⁻ pulse was observed in inoculated plots at T4 (September 2024), with burned plots showing a more modest and variable increase. Unburned soils maintained near-zero inorganic nitrogen concentrations throughout. (C) Metatranscriptomic profiles of nitrogen cycling genes at T0 and T5 across treatments. The heatmap shows normalized transcript abundance (log₂ CPM + 1) for key nitrogen cycling genes at T0 (November 2023) and T5 (December 2024) in burned, inoculated, and unburned plots (n = 3 per treatment at T0; n = 4 at T5). Ammonia monooxygenase subunit genes (amoA, amoB, amoC) are absent from burned soils at T0 but actively transcribed in unburned and inoculated soils. Denitrification and nitrite reduction genes (e.g., nirK, nirS, norB, nosZ, nrfA, nrfH, hao) show similar treatment-specific patterns. Partial recovery of nitrification transcripts is observed in burned plots by T5. Nitrogen fixation genes (nifH, nifK) were expressed at low levels across all conditions. Gene rows are grouped by biochemical pathway; each group is outlined with a black border.

In unburned soils, transcripts encoding ammonia monooxygenase subunits (*amoA*, *amoB*, *amoC*) were actively expressed. Metatranscriptomic profiling in burned soils at T0 revealed a near-complete absence of these transcripts, indicating an initial functional collapse of the first and rate-limiting step of nitrification (Fig. 4C). Inoculated plots showed immediate restoration of *amoA*/*B*/*C* and *hao* transcription at T0, demonstrating that the introduced community contained metabolically active ammonia-oxidizing organisms from the outset. Denitrification and dissimilatory nitrate reduction to ammonia (DNRA) transcripts (*nirS*, *nosZ*, *nrfA*, *nrfH*) were similarly depleted in burned soils relative to unburned soils at T0. However, denitrification and DNRA genes were transcribed in inoculated plots, indicating that inoculation re-established not only nitrification but a broader suite of nitrogen transformation pathways. Partial rebound of these transcript levels occurred in burn plots by T5. This suggests that natural succession gradually restored some N-cycling capacity to burned soils, although the effect on soil nitrogen species was smaller relative to inoculated soils. Nitrogen fixation (*nifH*) transcripts were low across all conditions and time points including unburned soils, indicating that N fixation plays a minor role in nitrogen inputs at this site, regardless of disturbance state.

To identify the specific organisms driving nitrification recovery, we tracked transcript levels of *amoA* mapping to MAGs at ≥ 99% ANI (Fig. 5A). In unburned plots, *amoA* reads mapped to sequences from six bacteria at T0 and to seven at the end of the experiment. Burned plots exhibited no *amoA* expression at T0, but transcripts mapped to two bacterial genomes at the last time point. In inoculated plots, *amoA* expression was attributed to seven bacterial genomes at T0; by the final timepoint, expression persisted in only two *Nitrosospira* (BFS-1554 and BFS-1557). These two persisting organisms were strains that also became active in burn plots at the end of the experiment. Across all timepoints, relative abundance data indicate that the two strains expressing *amoA* in the inoculated plots had relatively greater abundance at earlier timepoints relative to burned soils, and sustained this lead throughout the experimental period (Fig. 5B).

**Figure 5.**
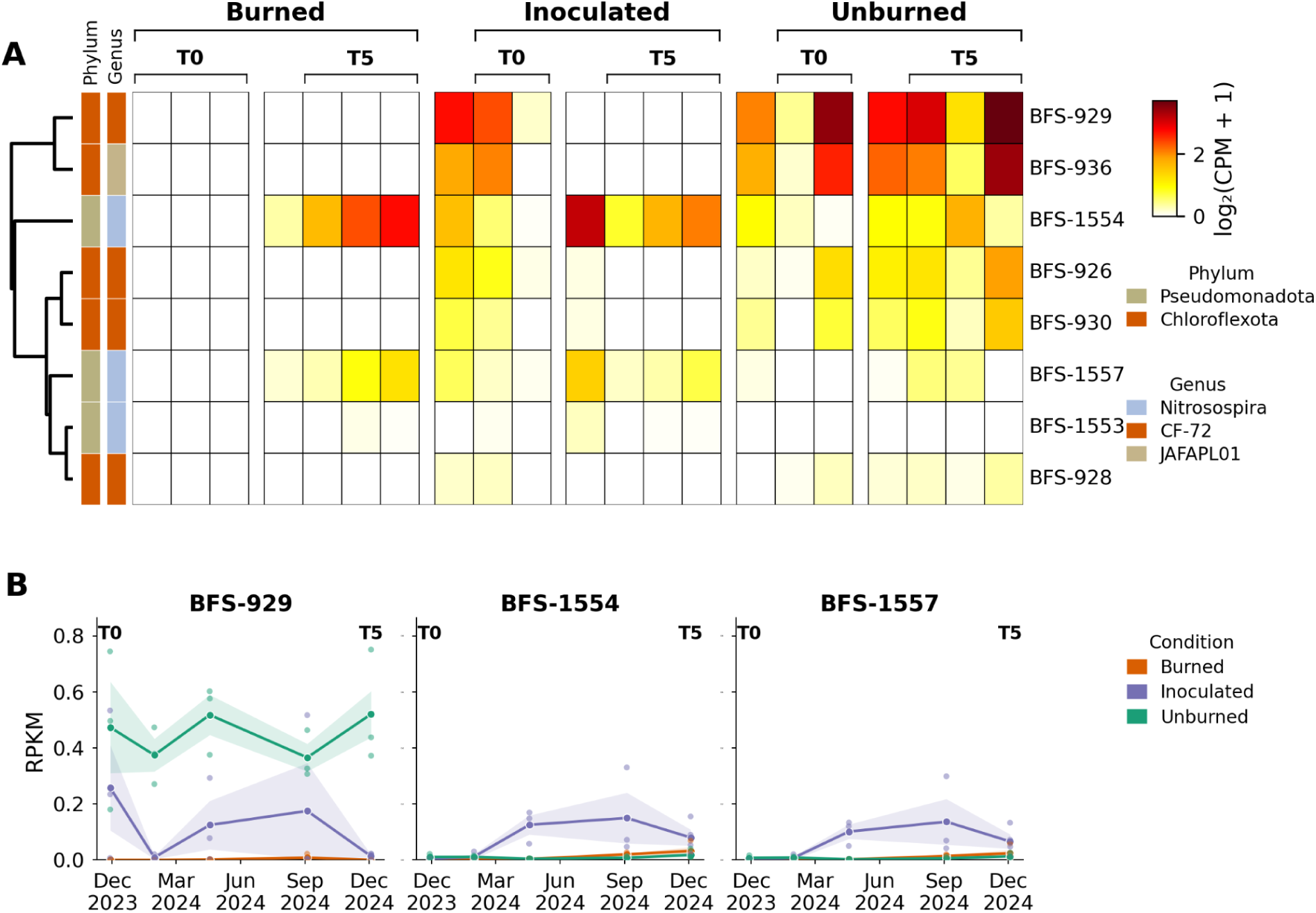
MAG-resolved ammonia monooxygenase (*amoA*) transcription and organism abundance across treatments and time at 99% ANI. (A) Transcription of *amoA* across metagenome-assembled genomes (MAGs) at 99% ANI for burned, inoculated, and unburned soils at T0 and T5. At 99% ANI, seven organisms expressed *amoA* in inoculated plots at T0 (CPM > 0 in ≥ 50% of replicates); by T5, only two persisted with active transcription across replicates. The surviving organisms (MAG IDs BFS-1554 and BFS-1557) independently emerged as the dominant nitrifiers in burn-only soils by T5. (B) Temporal abundance profiles (RPKM) of 3 of 7 amoA-expressing organisms (BFS-929, BFS-1554, and BFS-1557), plotted across all sampling time points for burned (red) and inoculated (blue) soils. Inoculated soils sustain higher *amoA*-transcribing biomass at earlier time points relative to burned controls, with seasonal fluctuations in abundance. Values in burned soils remain near-zero across most time points, with modest recovery at T4–T5.

An analysis of the activity of the enzyme hydroxylamine oxidoreductase (*hao*) that catalyzes the second step of nitrification revealed a similar pattern (Fig. S12). In unburned plots, *hao* transcripts mapped to six bacterial genomes. In the burned plots, no transcripts mapped to *hao* at the initial timepoint. In inoculated plots, transcripts mapped to five bacterial genomes at T0, although the activity was primarily detected in *Nitrosospira* BFS-1552 and 1554. By the end of the experiment, *hao* expression was detected for three organisms in inoculated plots (predominantly BFS-1552 and 1554) and for only BFS-1552 and 1554 in burned plots.

The strains BFS-1552, 1554 and 1557 of the genus *Nitrosospira* are predicted to have relatively slow growth rates (e.g., > 5h, which is the upper bound of gRodon’s reliable estimation) and predicted optimal pH for growth (pH 6.9 - 7.0), close to the pH of the burned and inoculated soils. In contrast, the predicted optimal pH for growth of the bacteria no longer active in inoculated plots at the final timepoint was lower (e.g., for the Chloroflexota BFS-929, BFS-936 that express *amoA*, predicted pH 5.6 and 5.4, respectively).

### Aromatic carbon catabolism is upregulated in post-fire soils

Metatranscriptomic analysis at the initial and final time points revealed condition-related differences in the expression of aromatic carbon degradation pathways (Fig. 6). Aromatic catabolism genes constituted a greater proportion of total transcriptional output in burned and inoculated soils than in unburned reference soils at T0. Catechol ortho-cleavage genes (*catA*, *catB*), protocatechuate ortho-cleavage genes, and catechol meta-cleavage genes were all more highly represented in burned and inoculated soils, whereas expression of the protocatechuate 4,5-cleavage genes (*ligA*, *ligB*, *ligC*, *ligI*, *ligJ*, *ligK*) was similar across all three conditions.

**Figure 6.**
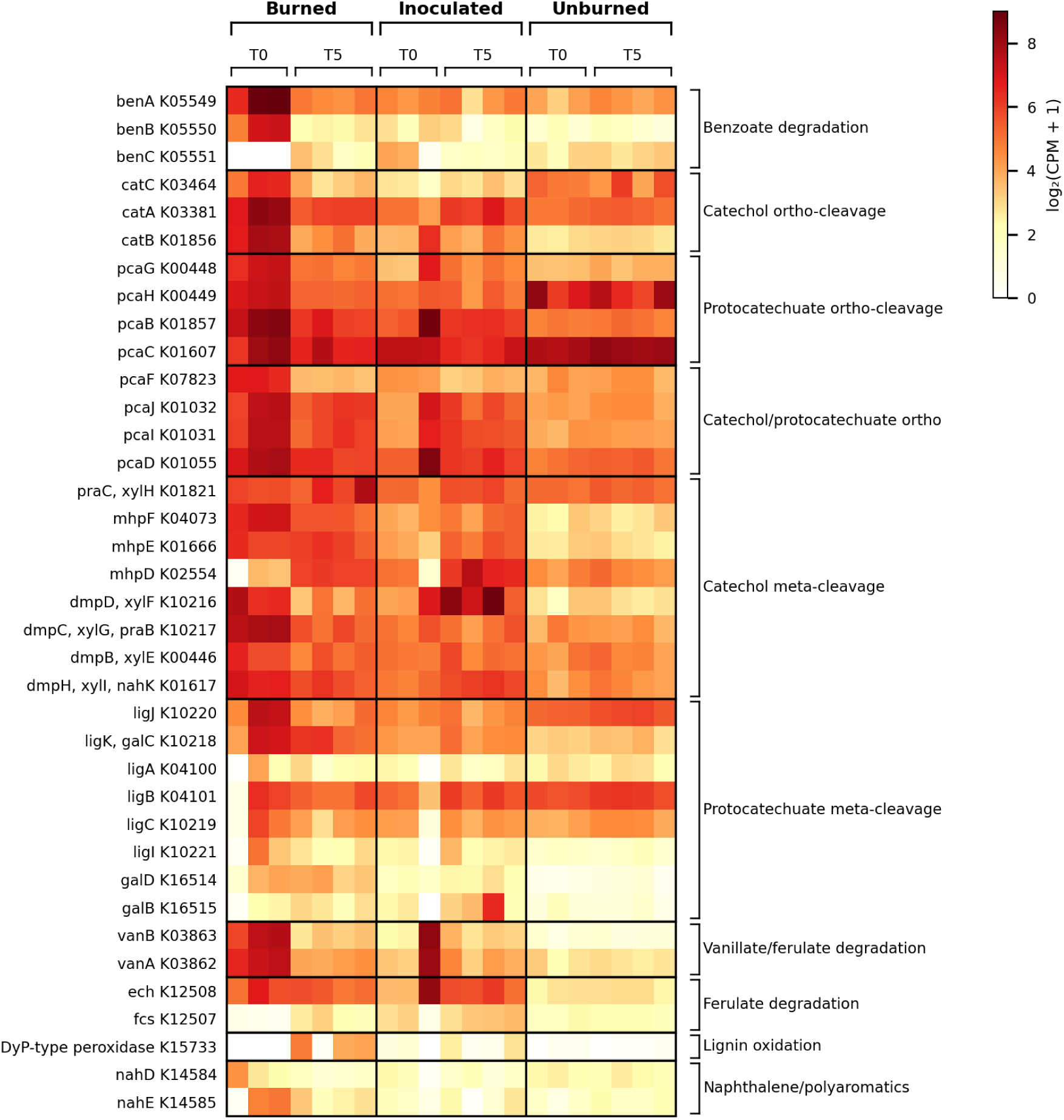
Metatranscriptomic profiles of aromatic carbon catabolism genes at T0 and T5 across treatments. Heatmap showing normalized transcript abundance of genes encoding aromatic carbon catabolism, including catechol ortho-cleavage, protocatechuate ortho-cleavage, and protocatechuate meta-cleavage, and benzoate, vanillate, ferulate, lignin and naphthalene degradation at T0 and T5 for burned, inoculated, and unburned plots. Burned and inoculated soils show greater proportional transcription of ortho-cleavage genes relative to unburned soils at T0, consistent with elevated pyrogenic aromatic substrate availability following combustion. Protocatechuate meta-cleavage gene expression was similar across all three conditions.

Burned soils harbored significantly fewer organisms transcribing *catA* at T0 compared to unburned soils across all taxa (Welch’s t-test, p < 0.05; Fig. S13), consistent with fire-induced loss of microbial diversity. Inoculated plots showed greater taxonomic richness of *catA*-expressing Actinomycetota and Pseudomonadota at T0 relative to burned-only plots. By T5, expression patterns in inoculated and burned-only soils had converged.

## Discussion

Following high-severity burns, the dominance of copiotrophic and stress-tolerant bacteria indicates that the post-fire surface community selects for fast-growing generalists able to exploit pyrogenic substrates under the altered chemistry of ash-laden soil, consistent with findings of prior studies^10,32^. The inoculant reintroduced fire-sensitive bacteria whose abundance recovered partially toward unburned levels, but the unburned community was not fully restored within the year. Notably, this clear compositional shift did not register as a difference in bulk alpha diversity likely due to high among-sample heterogeneity and the low abundances of introduced taxa. This reflects the fact that community-wide diversity indices can be insensitive to ecologically meaningful compositional change^29^. Ultimately, the persistently altered post-fire surface environment selected for the same bacteria in inoculated and burned plots.

As for bacteria, fungal diversity was reduced in burned compared to control plots. Fire drove a pronounced mycorrhizal-to-saprotrophic lifestyle shift representing the functional dismantling of the plant–fungal mutualistic network. Ectomycorrhizal fungi (EMF) are functionally obligate mutualists for most temperate conifers and are among the most dispersal-limited members of the forest soil microbiome; their loss following fire is widely considered a primary constraint on natural regeneration, particularly on large high-severity patches where spore dispersal from adjacent intact forest is increasingly distance limited^21,22^. Here, a single surface inoculation restored a diverse EMF assemblage that mirrored the surrounding intact forest. Simultaneous presence at the surface and at depth may reflect active hyphal extension through the soil profile, spore percolation with precipitation, or physical translocation during raking. EMF existence in the absence of mature conifer hosts suggests that introduced fungi subsisted on alternative carbon sources and/or fine root material carried in the inoculum. The co-restoration of EMF and ErM suggests that inoculation with soil provides a complex mycorrhizal functional spectrum.

This is a promising approach, the alternative being to use individually laboratory-cultivated fungal species as probiotics^33,34^. EMF and ErM establishment in inoculated plots was likely temporary due to depletion of resources and the absence of a symbiotic partner, but we cannot rule out the possibility that these fungi would have rebounded at a later timepoint. Nonetheless, inoculation provides a meaningful window during which restoration tree planting could take advantage of a pre-established mycorrhizal network. However, direct evidence that seedlings benefit from this residual inoculated EMF community is needed. The potential for restoration of a complex fungal microbiome is a key outcome of this study, with direct relevance to post-fire forest regeneration.

High-severity fire persistently reversed the microbial life-history trend at the soil surface, shifting communities from a K-selected state, in which slow-growing specialists dominate under stable, resource-limited conditions, to an r-selected state in which fast-growing opportunists dominated. While fast growth has been identified as a trait of post-fire bacterial colonizers through genome-resolved analysis^10^ and rRNA operon copy number^35^ approaches, our data demonstrate the shift as a sustained change throughout the initial year following a high-severity fire event. Subsurface communities (5–10 cm) recovered toward K-selection substantially faster than surface soils, likely reflecting the more buffered abiotic conditions at depth and consistent with prior observations that fire effects on microbial communities are attenuated with soil depth^4,10^. In inoculated soils, the dynamic and variable growth rate–abundance relationship is indicative of fluctuating r vs. K-selection. This may be governed by abiotic conditions (soil chemistry, carbon and nitrogen substrate availability, reduced moisture retention) that intermittently allow K-strategists to proliferate. Full restoration of K-selection may therefore require amelioration of the abiotic environment through organic matter recovery and vegetative return.

Incomplete compositional convergence between inoculated and unburned communities is likely attributable, in part, to ash-related elevated pH, which selected for organisms with higher pH growth optima. Thus, the post-fire soil environment is a selective force that cannot be immediately overcome by microbial reintroduction, either by natural dispersal or inoculation. Recovery of circumneutral pH via precipitation-mediated leaching, organic matter accumulation, and returning microbial acid production, may therefore be an integral component of microbiome recovery toward the unburned state, and a prerequisite for full compositional convergence.

Metatranscriptomic profiling and soil nutrient measurements captured nitrogen cycling differences across conditions. The near-complete absence of *amoA/B/C* transcripts in burned soils at T0 indicates functional silencing of nitrification by fire, consistent with the persistent ammonium accumulation we observed and with prior reports of reduced ammonia-oxidizer abundance and shifted ammonia-oxidizer community structure in burned mixed conifer soils^36^. A prior cross-system meta-analysis established that post-fire NH₄⁺ peaks within months and declines toward prefire levels by one year, while NO₃⁻ accumulates more slowly and peaks at 7–12 months post-fire^37^. Recent metagenomic work in chaparral documented a rise in nitrification gene abundance over the first year after fire^38^. We find that inoculation immediately restored transcription of *amoA/B/C*, denitrification and dissimilatory nitrate reduction genes, with correspondingly faster and greater NH₄⁺ drawdown and NO₃⁻ accumulation in inoculated compared to burned plots.

Genome-resolved tracking demonstrated that inoculation accelerates succession rather than installing a distinct community. In both inoculated and burn plots, two dominant nitrifying strains of *Nitrosospira* emerged, paralleling results from burned mixed conifer soils^36^. The *Nitrosospira* strains are predicted to have an optimal growth pH well suited to ash-laden post-fire soils and predicted slow growth rates that would make their establishment and activity lag behind fast-growing r-strategists in the absence of inoculation. Overall, our results suggest that inoculation’s contribution to the documented recovery of nitrogen cycling is the early seeding of slow-growing, environmentally matched specialists that would otherwise arrive only after extended dispersal-limited succession.

High-severity fire substantially increases the pool of aromatic carbon compounds in soil through incomplete combustion of biomass^39^. The proportional enrichment of aromatic catabolism transcripts in burned and inoculated soils is consistent with a community-level reorientation toward pyrogenic substrate utilization, corroborating findings from conifer forest burns in the Colorado and Wyoming Front Range^10^. Protocatechuate 4,5-cleavage expression, by contrast, was similar across all conditions, suggesting that this branch reflects an ongoing baseline process in conifer forest soils. The fire-associated upregulation of the catechol and protocatechuate ortho-cleavage routes, which converge on β-ketoadipate, feeding directly into the TCA cycle, instead points to a shift in substrate availability following combustion. The introduction of Actinomycetota and Pseudomonadota species via the inoculant added capacity for aromatic compound catabolism. By the final timepoint, inoculated plots showed slightly more richness of *catA* (the initial step in catechol catabolism) expressing organisms compared to burn plots. This pattern is consistent with the general conclusion that inoculation widens early functional diversity but ultimately there is selection for environmentally compatible introduced taxa.

This study utilized burn piles, which permitted experimental control over temperature and proximity to the inoculant source (thereby holding abiotic factors constant between conditions). However, burn piles differ from natural wildfires in spatial extent, combustion quantities, and burn timing (given that many natural fires occur after drought). Nonetheless, our results stem from field experiments that, although somewhat limited by substantial natural soil heterogeneity and small replicate size, capture processes that occurred under near-realistic conditions that could not be approximated in laboratory experiments. Whether the inoculation effects documented here generalize to large high-severity natural wildfires remains to be established.

Although the findings of this study demonstrate the benefit of direct soil amendment, it may be worth considering augmentation by targeted amendment of certain cultivable organisms that our research suggests may be important for soil microbiome recovery. For example, we identified specific slow-growing nitrifiers, EMF fungi, and aromatic carbon degraders with high post-fire persistence, suggesting that these organisms are capable of establishing in the hostile burned soil environment. Introduction of early succession organisms, either by unburned soil amendment or culture-based inoculation, opens niches for other microbiome members.

An important consideration related to deployment of the soil inoculation approach is how to implement the strategy cost effectively and at the scale of wildfire burn scars. It is unlikely to be feasible to comprehensively distribute unburned soil over tens to hundreds of square kilometers. It is established that wind-based microbial dispersal is important in natural recovery of soil from disturbance^20^, but such inputs are often slow and decline with increasing distance. Thus, we anticipate an effective implementation could involve creation of inoculated patches from which dispersal can proceed. In the current study, the unburned soil was raked into burned soil, but it remains unclear whether this is key to microbial engraftment. If raking is important, we suggest very shallow mechanical raking, such as used in agricultural tine-based seeding. Alternative options could involve aerial dispersal of soil from e.g., drones, perhaps during rain events to improve penetration into soil. The appropriate spacing of local inoculation sites to optimize dispersal may be predicted based on topography, dominant wind direction, hydrological-flow pathways, erosion risk, and burn severity patterns. For example, inoculation sites at ridge tops rather than on the valley floor may improve the probability of dissemination of soil particles and microbes by wind and water flow.

Conceptually, bulk soil inoculation is an attractive strategy because it delivers a broad suite of microbial functions simultaneously, including nitrifiers, mycorrhizal fungi, and aromatic carbon degraders. In principle, repeated applications of native soil over time could further enhance establishment by allowing different phases of the recovering community, including taxa that fail to persist after a single application, to colonize under changing post-fire conditions. An appealing attribute of the unburned soil inoculation approach is cost: it is “cheap as dirt”.

## Conclusions

Our results demonstrate that a single inoculation of unburned soil into high-severity burn sites accelerates the recovery of prokaryotic taxa, mycorrhizal fungi, and nitrogen cycling activity across a 12-month post-fire period. Yet a consistent finding across all analytical layers (prokaryotic diversity, fungal composition, genome-resolved life-history strategies, and metatranscriptomic functional profiles) is that bulk soil inoculation acts as an accelerant of environmentally determined succession rather than as a stable community transplant. The organisms that ultimately dominate nitrogen cycling, aromatic carbon catabolism, and the surface bacterial community are those selected by the post-fire abiotic environment; inoculation’s contribution is to ensure that functionally important but slow-colonizing taxa are present and active during the early post-fire window rather than arriving only after years of dispersal-limited succession. This distinction between temporal acceleration and community override has direct implications for the design and evaluation of microbiome-based restoration. As wildfire extent and severity continue to expand under climate change, and natural regeneration increasingly fails on large burned patches, scalable biological tools for collapsing the timeline of soil microbiome recovery warrant further investment.

## Methods

### Study site and experimental design

The study was conducted at the University of California, Berkeley’s Blodgett Forest Research Station (BFRS) in the central Sierra Nevada Mountains (1,300 m elevation; 38.9119° N, 120.6693° W). Vegetation at BFRS is dominated by a mixed conifer forest type composed of variable proportions of white fir (*Abies concolor*), incense-cedar (*Calocedrus decurrens*), Douglas-fir (*Pseudotsuga menziesii*), sugar pine (*Pinus lambertiana*), ponderosa pine (*Pinus ponderosa*), and California black oak (*Quercus kelloggii*)^40^.

Burn piles were constructed within the canopy of the mixed conifer forest using locally sourced logs representing the dominant tree species. The burn pile used in this study measured approximately 6.4 × 6.2 × 2.1 m. The pile was ignited and allowed to burn and smolder to natural extinction over approximately two weeks. Soil surface temperatures were recorded continuously throughout the burn using type K thermocouple probes (Part # HH-K-20-25) affixed to the soil surface within the burn and connected to HOBO 4-Channel Thermocouple Data Loggers (Part # UX120-014M) in Onset Protective enclosures (Part # CASE-4X-2) wrapped in insulating mineral wool and wildland fire shelters, which were buried external to the burn site.

The burn pile footprint was subdivided into a grid of 1 × 1 m subplots. Subplots were assigned in an alternating checkerboard pattern to one of two conditions: untreated burn control (n = 3) or soil inoculation (n = 3). An additional three unburned subplots were established in the surrounding intact mixed conifer forest approximately 10 meters from the perimeter of the burn pile. Soil used for inoculation was collected from 0–10 cm depth at 10 randomly selected locations adjacent to the unburned plots, pooled into buckets, and manually mixed to homogenize. Inoculation was performed approximately two weeks after the fire naturally extinguished. The pooled inoculant was lightly raked into the upper 0–5 cm of the inoculated subplots. Burn control subplots were similarly raked to 0–5 cm depth to control for any physical disturbance of the soil surface.

### Soil sampling and processing

Soils were collected at six time points: T0 (November 28th, 2023), T1 (December 14th, 2023), T2 (January 30th, 2024), T3 (April 18th, 2024), T4 (September 4th, 2024), and T5 (December 4th, 2024), spanning a full annual cycle encompassing winter and summer. At each time point, two independent soil samples were collected from each subplot: one for molecular (–omics) analyses and one for soil chemistry.

For molecular analyses, a single soil core (2” diameter) was collected from each subplot using a soil corer and subdivided in the field into 0–5 cm and 5–10 cm depth fractions. Sample fractions were immediately sealed in sterile bags and flash-frozen in liquid nitrogen in the field. Frozen samples were transported to the laboratory on dry ice and stored at –80°C until processing, which occurred within six months of collection. For soil chemistry, a separate sample was collected from each subplot at 0–5 and 5–10 cm depths using a hand trowel. Chemistry samples were stored on ice during transport and processed upon return to the laboratory.

Each subplot was sampled once per time point, yielding n = 3 samples per condition (burned, inoculated, unburned) at T0–T4, and n = 4 samples per condition at the final time point T5.

### Soil chemistry

Soil pH and moisture levels were measured within 24 hours of collection. Soil pH was measured in a 1:2 soil:water suspension (10 g fresh field-moist soil in 20 mL deionized water), allowed to equilibrate for 30 minutes, and measured with a glass electrode pH meter calibrated against pH 4, 7, and 10 standard buffer solutions. Gravimetric soil moisture content (MCd) was determined by drying ∼10 g of fresh field-moist soil in pre-weighed aluminum tins at 105°C to constant mass (≥24 h, with repeat weighings until consecutive measurements differed by <0.001 g). Moisture content was calculated as the mass of water lost divided by oven-dry soil mass. Soil NO₃⁻ and NH₄⁺ were extracted immediately upon return to the laboratory by shaking 5 g field-moist soil in 50 mL 2M KCl for 1 hour, settling, and filtering through Whatman filter paper. Filtered extracts were frozen at −20°C and shipped on dry ice to Ward Laboratories, Inc. (Kearney, NE) for quantification by flow injection analysis (FIA). Remaining soil was dried in paper bags at ∼45°C prior to shipping to Ward Laboratories, Inc. for additional nutrient analyses. Soil organic matter was determined by loss on ignition (LOI; 360°C, 2 h following desiccation at 105°C). Available phosphorus was extracted with Mehlich 3 solution; exchangeable cations (K, Ca, Mg, Na) with ammonium acetate (pH 7.0); and micronutrients (Zn, Fe, Mn, Cu, B) with DTPA-sorbitol (pH 7.3); all quantified by inductively coupled plasma (ICP). Chloride was extracted with calcium nitrate solution and measured by FIA. Total soil carbon and nitrogen were measured by combustion. Differences in soil chemistry between conditions, timepoints, and depths were determined using two-sample t-tests assuming equal variances.

### DNA extraction, 16S rRNA gene and ITS amplicon sequencing

Total nucleic acids were co-extracted from 1.5–2 g of homogenized soil per sample (targeting 1.25 g dry-weight equivalent) using the RNeasy PowerSoil Total RNA Kit (QIAGEN, cat. no. 12866-25), with DNA recovered from the same lysate via the RNeasy PowerSoil DNA Elution Kit (QIAGEN, cat. no. 12867-25), following the manufacturer’s instructions. Co-extraction ensured that DNA and RNA fractions used for downstream amplicon, metagenomic, and metatranscriptomic analyses originated from identical sample material. DNA concentration was quantified by Qubit fluorometry (Thermo Fisher Scientific), and extracts were stored at −80 °C until library preparation.

Bacterial and archaeal communities were characterized by amplicon sequencing of the 16S rRNA gene V4 region using the primers 515F/806R; fungal communities were characterized by amplicon sequencing of the ITS1 region using the primers ITS1F and ITS2. Library preparation and sequencing for both marker genes were performed by Novogene Corporation on the Illumina NovaSeq 6000 platform (250 bp paired-end), with quality filtering to Q30 ≥ 75%. Each marker gene was sequenced across two batches.

Raw reads were processed in the QIIME2 environment (release 2024.5)^41^. DADA2^42^ was used to filter reads, learn error rates, denoise, and remove chimeras, producing amplicon sequence variants (ASVs) for each marker. Taxonomy was assigned to bacterial and archaeal ASVs using the QIIME2 sklearn classifier trained on the SILVA 138 database (99% sequence similarity), and to fungal ASVs using the sklearn classifier trained on the UNITE fungal ITS database (v10.0, release 2024-04-04, 99% clustering). Feature tables, representative sequences, and taxonomy assignments were merged across batches at the QIIME2 artifact level prior to downstream analysis.

After DADA2 filtering, 16S samples retained 91,259–247,892 reads (mean 161,479; n = 117), and ITS samples retained 62,586–273,084 reads (mean 154,620). For alpha diversity analyses, feature tables were rarefied to 91,000 reads per sample for 16S and 62,500 reads per sample for ITS (depths chosen to retain all samples) and Shannon diversity was calculated using the QIIME2 diversity plugin. Differences in Shannon diversity between conditions at each timepoint and depth were tested using one-way ANOVA with Bonferroni-corrected pairwise t-tests.

Ecological lifestyles were assigned to fungal ASVs using FUNGuild (v1.1)^43^. Following FUNGuild creator recommendations, only assignments with a Confidence Ranking of “Probable” or “Highly Probable” were retained for downstream analyses. Guild-level relative abundance was calculated as the proportion of total fungal reads assigned to each guild per sample, and EMF genus richness was calculated as the number of unique EMF genera detected per subplot across all retained ASVs.

Bray-Curtis dissimilarity matrices were generated from rarefied 16S and ITS feature tables in QIIME2 and exported to R (v4.5.1) for downstream analysis using the vegan^44^ package (v2.8.0). Non-metric multidimensional scaling (NMDS) ordinations were performed with metaMDS (k = 2,100 random starts), separately for surface (0–5 cm) and subsurface (5–10 cm) samples.

Condition effects on community composition were tested by permutational multivariate analysis of variance (PERMANOVA; adonis2, formula = Bray-Curtis ∼ condition × timepoint, 9,999 permutations, by = “terms”)^45^. Multivariate homogeneity of group dispersions was assessed using betadisper with permutest (9,999 permutations) and Tukey’s HSD for pairwise comparisons^46^.

### Metagenomic assembly and binning

The same DNA extracted for 16S rRNA amplicon sequencing was used for metagenomic sequencing. Samples from timepoint T1 were excluded from metagenomic sequencing due to the short interval between T0 and T1 (∼2 weeks) and project budget constraints. Metagenomic libraries were prepared and sequenced by Novogene Corporation on the Illumina NovaSeq X Plus platform (150 bp paired-end) across two batches (n = 39 and n = 60 samples; 99 samples total), targeting 30 Gb of raw sequence per sample with quality filtering to Q30 ≥ 85%.

Raw reads were processed using BBTools to trim Illumina adapters and remove PhiX174 spike-in and Illumina artifact contaminants, followed by quality trimming with Sickle (paired-end mode, Sanger encoding, quality threshold Q20, minimum length 20 bp). Read quality was inspected before and after processing using FastQC.

Quality-controlled reads from the three replicate subplots per condition × depth × timepoint group were pooled prior to co-assembly with MEGAHIT v1.2.9^47^ with k-mer lengths 27–127 (step 10) and minimum contig length 1,000 bp. Per-group co-assembly was chosen over per-sample assembly to improve contiguity and recovery of low-abundance taxa shared across replicate subplots while preserving condition- and timepoint-specific community signals. Open reading frames on contigs ≥ 1,000 bp were predicted with Prodigal v2.6.3^48^ in metagenomic mode (-p meta -m).

Metagenome-assembled genomes (MAGs) were recovered from each co-assembly using the Banfield-lab ggBin pipeline, which runs CONCOCT, MetaBAT2, and VAMB in parallel and integrates their outputs with DAS Tool^49^ for ensemble bin refinement. Bin completeness and contamination were estimated using CheckM v1.2.2^50^ with the “lineage_wf” workflow. Bins meeting MIMAG draft-quality criteria (≥ 50% completeness, ≤ 10% contamination)^51^ were retained for analysis.

MAGs were dereplicated using dRep^52^ at two ANI thresholds, 95% (species-level clustering) and 99% (strain-level clustering). The 95% ANI set was used for community-level growth-rate and life-history analyses, while the 99% ANI set was used for strain-resolved functional analyses and metatranscriptome read mapping. MAG taxonomy was assigned with GTDB-Tk v2.4.0 using the “classify_wf” function with the GTDB R220 release. Quality-controlled reads from all 99 samples were mapped to each dereplicated MAG set using Bowtie2 v2.5.4^53^. Genome-level relative abundance, mean and trimmed-mean coverage, and per-MAG read counts were calculated using CoverM v0.7.0^54^ in genome mode.

### MAG annotation and trait inference

Predicted open reading frames were functionally annotated by performing protein similarity searches against the KEGG (release 2015_06), UniRef100 (release 2014_02), and UniProt (release 2015_06) databases using DIAMOND blastp (v2.1.10)^55^ in sensitive mode. For pathway-level functional analyses, a curated set of KEGG Orthology (KO) identifiers derived from Nelson et al.^10^ was used to identify genes involved in nitrogen cycling and aromatic carbon catabolism, including ammonia oxidation, denitrification, DNRA, nitrogen fixation, and aromatic ring-cleavage pathways. For richness analysis of organisms with detectable *catA* transcription, MAGs were considered actively transcribing *catA* within a condition × timepoint group if CPM > 0 in ≥50% of replicates. The number of unique *catA*-transcribing organisms per sample was compared between conditions at T0 and T5 using Welch’s two-sample t-tests. Maximum doubling times were estimated via codon usage bias using gRodon^30^ (v2.5.2); Spearman correlations between estimated doubling time and per-MAG relative abundance were computed within each condition × depth × timepoint group, with 95% confidence intervals estimated via bootstrap resampling (1,000 iterations, percentile method). Optimal pH for growth was predicted for each organism using a machine learning model trained on profiles of 56 genes encoded within the MAGs^56^. Predicted optimal pH was compared between conditions within each timepoint × depth group using one-way ANOVA, with Tukey-adjusted pairwise contrasts of estimated marginal means.

### Metatranscriptomic sequencing and quality control

Total RNA was extracted from 1.5 - 2 g of frozen soil from surface samples based on dry weight equivalent to 1.5 g (0–5 cm depth) collected at two timepoints (T0: November 28, 2023; T5: December 4, 2024) across three conditions (burned, inoculated, and unburned; n = 7 per condition, 21 samples total) using the RNeasy PowerSoil Total RNA Kit (Qiagen). RNA and DNA were co-eluted from the same extractions. Where RNA concentrations from individual extractions were below the sequencing provider’s quality threshold as determined by Qubit fluorometry and Bioanalyzer, multiple extraction replicates from the same sample were pooled at the column binding step prior to elution to achieve sufficient input material. Libraries were prepared and sequenced on an Illumina NovaSeq X Plus platform as 150 bp paired-end reads, generating a mean of 192.6 ± 23.8 million read pairs per sample (range 166.0–270.9 million).

Raw reads were quality-trimmed using fastp (v0.23) with adapter autodetection, a minimum read length of 50 bp, and a minimum Phred quality score of 20, retaining a mean of 98.5% of reads (mean Q30 rate: 95.6%). Ribosomal RNA was removed by mapping trimmed reads against a combined rRNA reference database (SILVA, 32,762 clustered representatives) using BBMap (v39.10) with default sensitivity parameters. Ribosomal RNA comprised a mean of 1.3 ± 3.5% of reads per sample (range 0.1–14.8%), reflecting effective mRNA enrichment during library preparation.

### Metatranscriptomic read mapping, de novo assembly, and quantification

For community-level functional profiling, non-rRNA reads from each condition were pooled across samples and digitally normalized using BBNorm prior to co-assembly with rnaSPAdes (v3.15, k-mer sizes 47, 67, 87). Separate co-assemblies were performed for each condition, yielding 5.2 million contigs (N50 = 875 bp) for burned, 15.1 million contigs (N50 = 493 bp) for inoculated, and 28.2 million contigs (N50 = 381 bp) for unburned soils, reflecting the greater transcriptional diversity of the unburned community. Genes were predicted on each assembly using Prodigal (v2.6.3, meta mode) and annotated with eggNOG-mapper (v2.1.13, diamond mode, eggNOG v5.0.2 database). Individual sample reads were mapped to their respective condition co-assembly using Bowtie2 (v2.5.4, --very-sensitive-local) and read counts per gene were quantified using featureCounts (Subread v2.0, -t CDS -g ID -p --countReadPairs, unstranded). Genes were filtered to retain those with ≥ 5 counts in ≥ 2 samples. Expression was normalized to counts per million (CPM) using total reads mapped to that condition’s assembly as the library size denominator, and KEGG Ortholog (KO)-level expression was calculated by summing CPM values across all genes assigned to each KO.

For MAG-resolved gene expression profiling, non-rRNA reads from all 21 samples were mapped against the dereplicated 99% ANI MAG reference (1,952 MAGs, 1,540,707 contigs, 8,566,919 predicted genes) using Bowtie2 (--very-sensitive-local --no-unal), retaining only concordantly mapped read pairs. A mean of 63.7 ± 18.5% of reads mapped to the MAG reference (range 33.2–88.5%), with lower mapping rates in unburned samples consistent with the greater organismal diversity of undisturbed soil communities. Read counts per gene were quantified using featureCounts (-t CDS -g ID -p --countReadPairs, unstranded) and MAG genes were annotated using eggNOG-mapper (v2.1.13, diamond mode). Expression was normalized to CPM using total reads mapped to the complete MAG reference as the library size denominator. Gene-level counts were aggregated to the MAG level by summing across all genes assigned to each KO within a given MAG, and expression values are reported as log₂(CPM + 1).

## Acknowledgments

We thank Robert A. York for assistance with site selection, burn pile construction, and ignition at the Blodgett Forest Research Station. Additionally, we thank Kaitlin Creamer, Allison Guitor, Leylen Miloslavich, and Kaden DiMarco for assistance with field sampling, and Rohan Sachdeva for development and maintenance of the Banfield laboratory annotation pipeline. This research was supported by the Lab to Land Institute with funding from Chris Kohlhardt, the Fifty Years’ (50Y) Manifest Grants program (Grant no. 057085-001), and the Chan Zuckerberg Initiative (CZI; Grant no. 053910-001).

## Author Contributions

E.L.W. and J.F.B. designed the study. E.L.W. conducted fieldwork, sample processing, and data analysis. E.L.W. and J.F.B. wrote the manuscript.

## Data Availability Statement

Raw sequencing reads (16S rRNA, ITS, metagenomic, and metatranscriptomic) have been deposited at NCBI under BioProject PRJNA1477628.

## Supplementary Information

Figs. S1 to S13 Table S1

