## Supplementary Figures for "Microbial inoculation accelerates post-fire soil recovery in a mixed conifer forest"

### Supplementary Figures for
#### Microbial inoculation accelerates post-fire soil recovery in a mixed conifer forest

Elliot L. Weiss[1] and Jillian F. Banfield[1]*

[1]Department of Earth and Planetary Science, University of California, Berkeley, CA 94720, USA

A B C

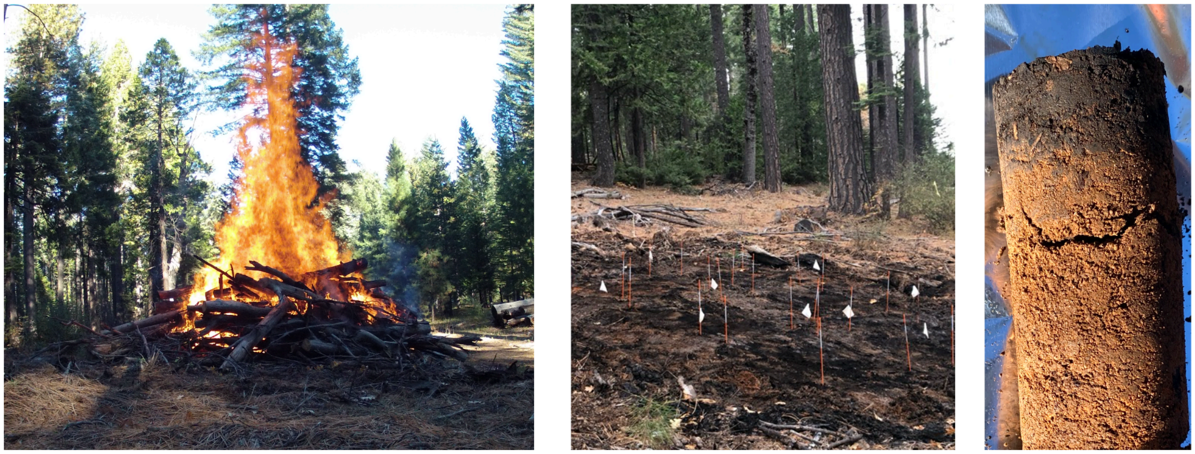

**Figure S1**. Experimental field site and burn pile in the Blodgett Forest. (A) High-severity experimental burn pile during peak combustion, constructed from locally sourced mixed conifer logs and ignited within the intact forest canopy. Peak soil surface temperatures reached approximately 800°C. (B) Burn site after natural extinguishment, showing the gridded subplot layout. Flagged stakes mark the 1 × 1 m subplots assigned to either burned or inoculated conditions. (C) Representative soil core from a burned subplot at time of inoculation, showing the dark ash-laden surface horizon transitioning to lighter mineral soil below.

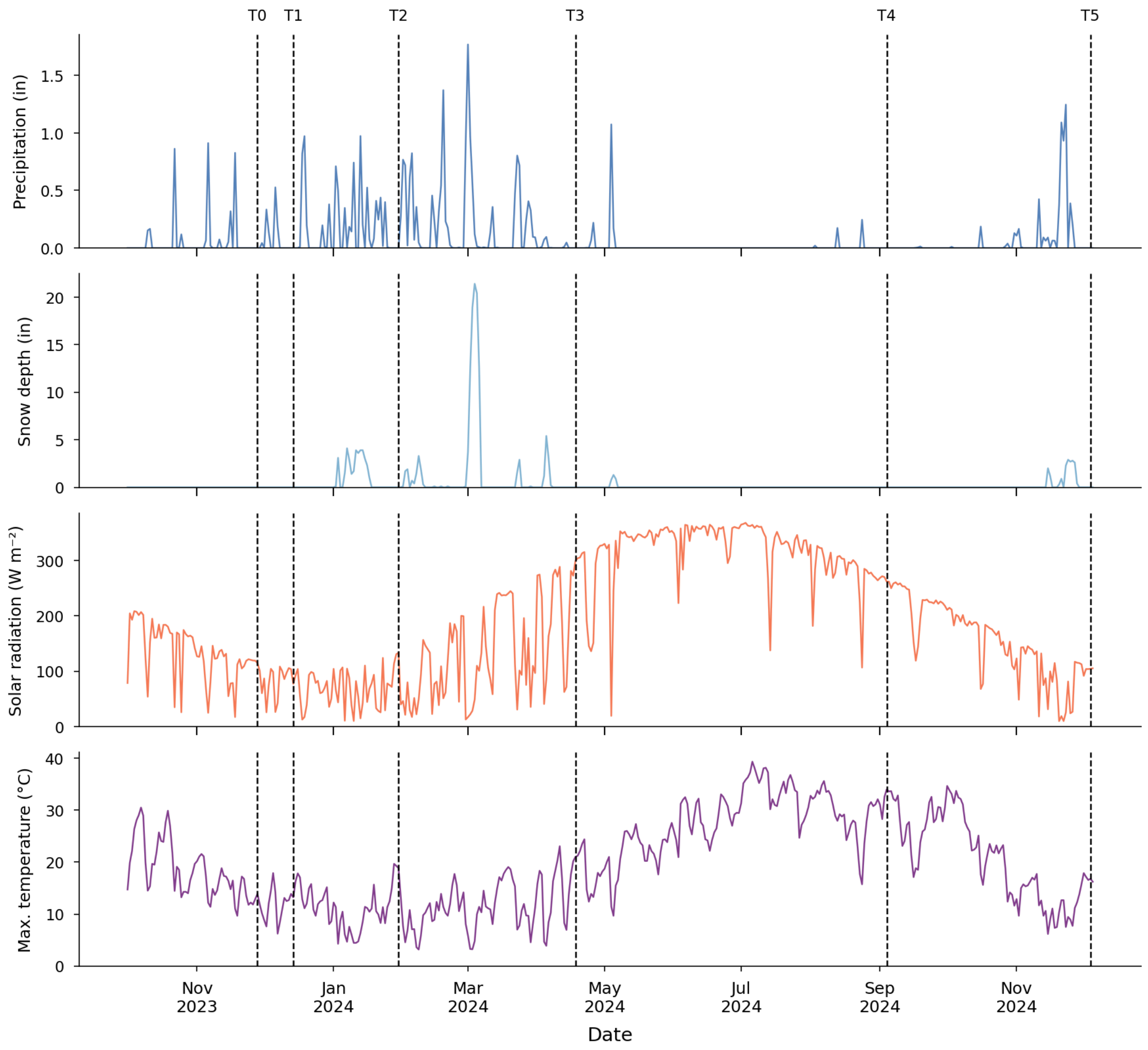


**Figure S2.** Climatological conditions and sampling timeline at Blodgett Forest Research Station, November 2023–December 2024. Time series of precipitation (inches), snowpack depth (inches), solar radiation (W $m^{-2}$), and air temperature (°F) recorded near the study site over the 12-month sampling period. Dotted red vertical lines indicate the six sampling time points (T0–T5). Snowpack depth remained moderate relative to preceding years, permitting consistent site access across all sampling events. The full annual hydrological cycle, including wet winters and a dry summer, is captured, providing climatically realistic context for interpreting community dynamics across treatments.

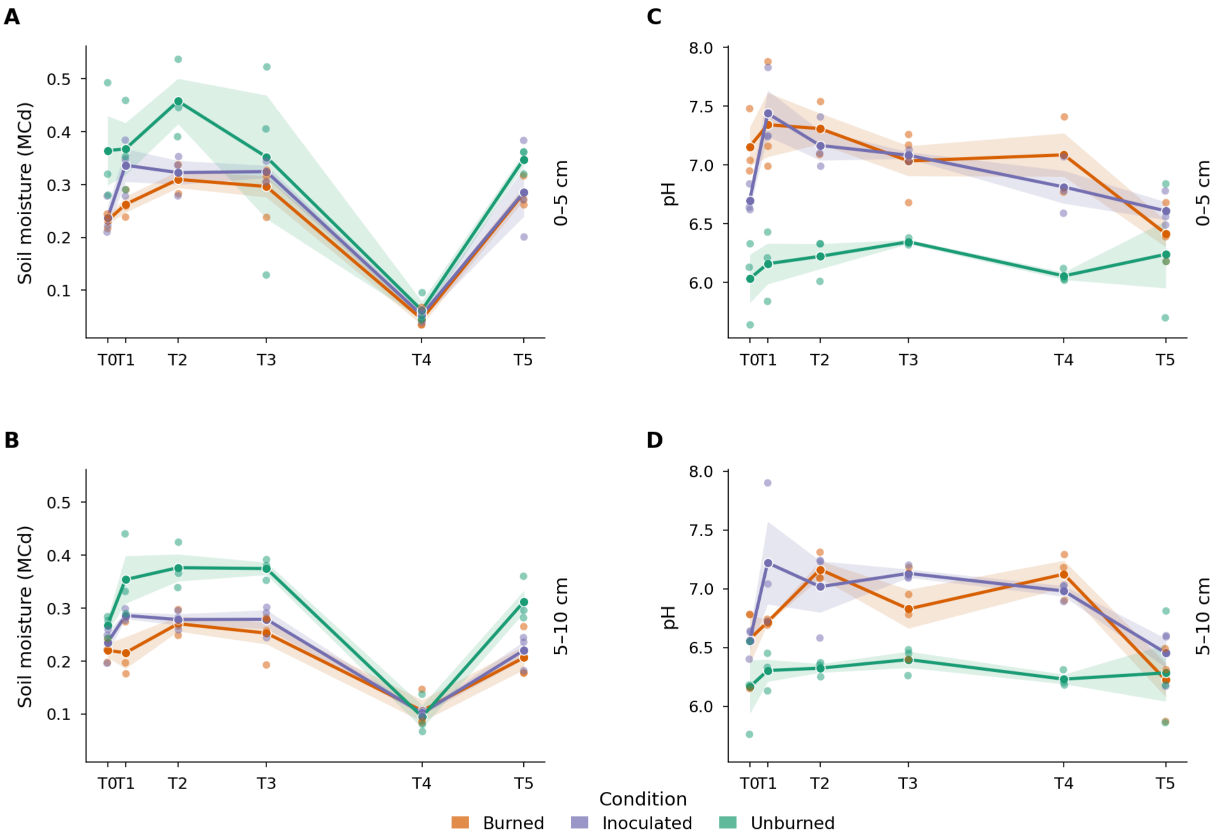


**Figure S3.** Soil moisture and pH dynamics in burned, inoculated, and unburned plots over the 12-month study period. Soil moisture (volumetric water content; MCd) at 0–5 cm (A) and 5–10 cm (B) depth, and soil pH at 0–5 cm (C) and 5–10 cm (D) depth, across all sampling time points (T0–T5). Lines connect treatment means; shaded bands indicate ±1 standard error; individual points show per-subplot values ($n = 3$ per treatment at T0–T4, $n = 4$ at T5). Unburned soils retained consistently higher moisture than burned and inoculated plots at both depths throughout the year, with the exception of the September 2024 (T4) sampling, when extended summer drought reduced moisture across all treatments. At 0–5 cm, burned plots exhibited significantly elevated pH at T0 relative to unburned soils (7.16 ± 0.28 vs. 6.03 ± 0.36; two-sample t-test, $t = 4.28$, $p = 0.013$), consistent with ash deposition following combustion. Inoculated plots also showed elevated pH at T0 relative to unburned soils (6.70 ± 0.12), reflecting ash inputs shared across the burn footprint. pH declined progressively in burned and inoculated plots across the sampling period, with both treatments approaching unburned values by T5 (burned: 6.42 ± 0.25; inoculated: 6.61 ± 0.15; unburned: 6.24 ± 0.57), likely reflecting a combination of precipitation-mediated leaching, returning microbial organic acid production, and plant litter inputs. At 5–10 cm depth, pH did not differ significantly between burned and unburned soils at T0 (6.57 ± 0.36 vs. 6.16 ± 0.40; two-sample t-test, $t = 1.31$, $p = 0.26$), suggesting that alkalinization effects were largely confined to the surface horizon at the time of initial sampling. Subsurface pH in burned and inoculated plots showed a delayed elevation peaking at T2–T3 before returning toward unburned values by T5.

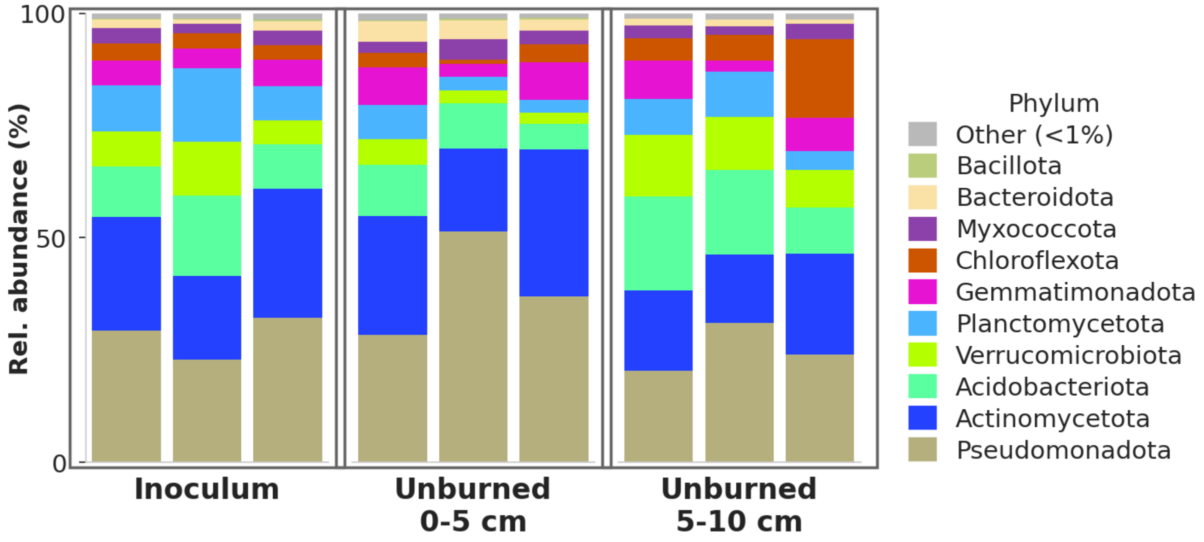


**Figure S4**. 16S rRNA relative abundance of inoculum in comparison to unburned samples at 0-5 and 5-10 cm at timepoint 0 (n = 3). Relative proportions of phyla in the inoculum used in this study are similar to and representative of unburned soils.

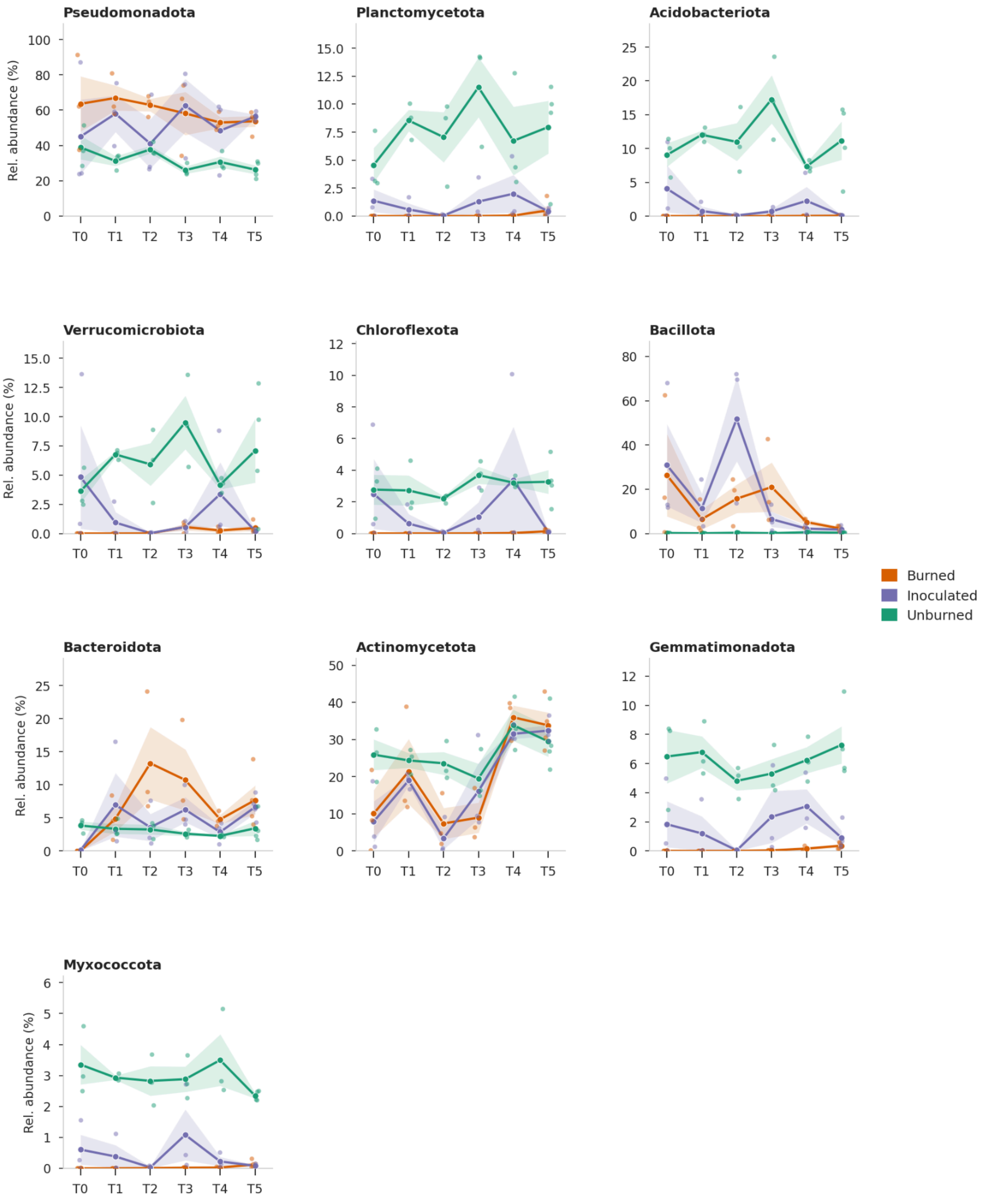
Pseudomonadota
Planctomycetota
Acidobacteriota
Verrucomicrobiota
Chloroflexota
Bacillota
Bacteroidota
Actinomycetota
Gemmatimonadota
Myxococcota
Rel. abundance (%)
T0
T1
T2
T3
T4
T5
Burned
Inoculated
Unburned

**Figure S5.** Relative abundance of major bacterial phyla across treatments and time points at 0–5 cm depth, estimated from 16S rRNA amplicon sequencing. Lines connect mean relative abundance values for each treatment at each time point; shaded bands indicate ±1 standard error; individual points show per-subplot values (n = 3 per treatment at T0–T4, n = 4 at T5). At T0, burned soils were dominated by Pseudomonadota, Actinomycetota, and Bacillota, characteristic of early post-fire copiotrophic succession, with minority phyla (Planctomycetota, Acidobacteriota, Verrucomicrobiota, Chloroflexota, Gemmatimonadota, Myxococcota) at or near zero. Inoculated plots showed substantially higher relative abundance of minority phyla from T0 onward, with Acidobacteriota, Verrucomicrobiota, and Chloroflexota recovering toward levels observed in unburned soils over the sampling period. Unburned soils maintained a compositionally diverse community throughout, with no single phylum dominating to the extent observed in burned plots.

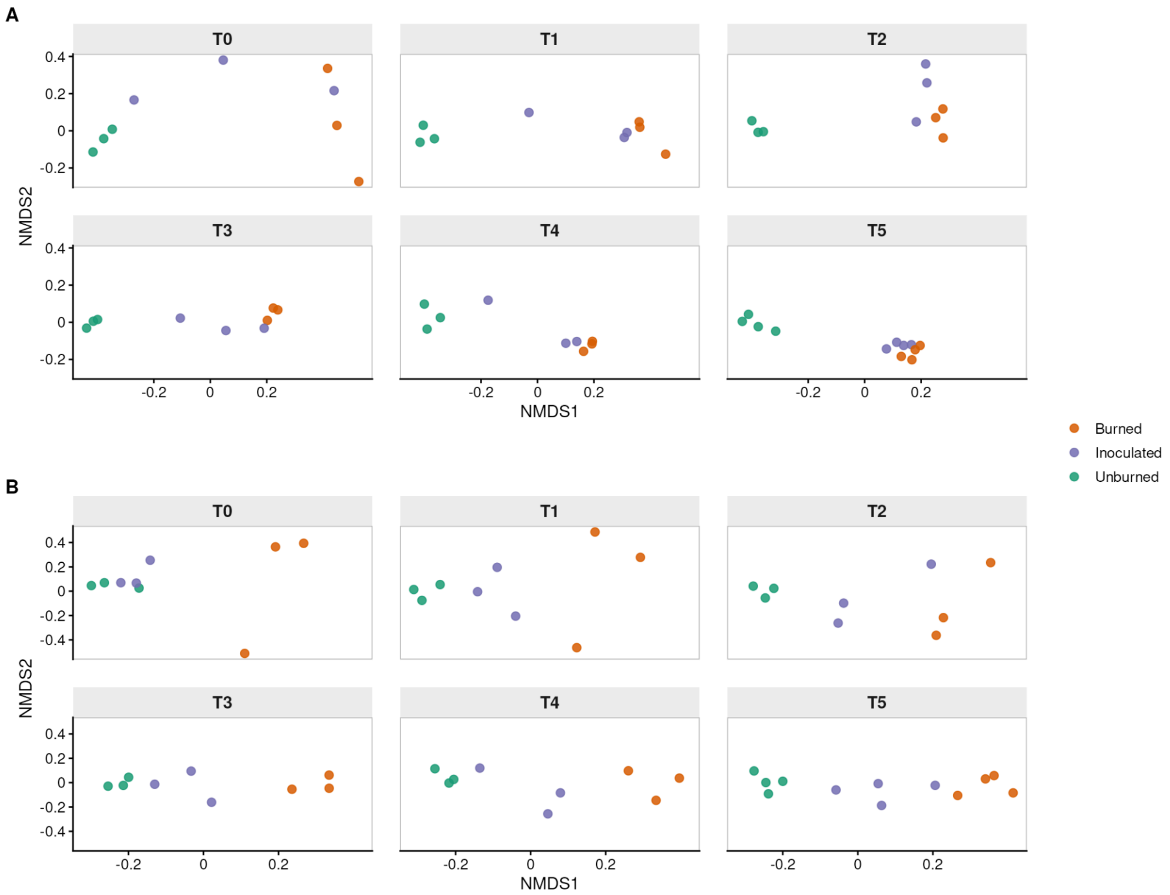


**Figure S6.** Non-metric multidimensional scaling (NMDS) of Bray–Curtis dissimilarity for bacterial (A) and fungal (B) communities at 0–5 cm depth across treatments and time points. Each point represents one subplot replicate (n = 3 per treatment at T0–T4, n = 4 at T5), colored by treatment. Panels show per-timepoint ordinations to facilitate visualization of community trajectories. (A) Bacterial community composition based on 16S rRNA amplicon sequencing (NMDS stress = 0.10; PERMANOVA $R^2 = 0.65$, $F = 4.30$, $p = 0.001$). Burned and inoculated plots exhibited significantly greater among-sample dispersion than unburned soils (PERMDISP, $F = 16.5$, $p = 2.5 \times 10^{-6}$; Tukey HSD $p < 0.001$ for both comparisons), while dispersion did not differ between burned and inoculated plots ($p = 0.42$). (B) Fungal community composition based on ITS amplicon sequencing. Treatment-level separation was maintained throughout the study period (NMDS stress ≈ 0.10; PERMANOVA $R^2 = 0.45$, $F = 1.91$, $p = 0.001$). Burned and inoculated plots were more dispersed than unburned soils (PERMDISP Tukey HSD $p < 0.001$ for both).

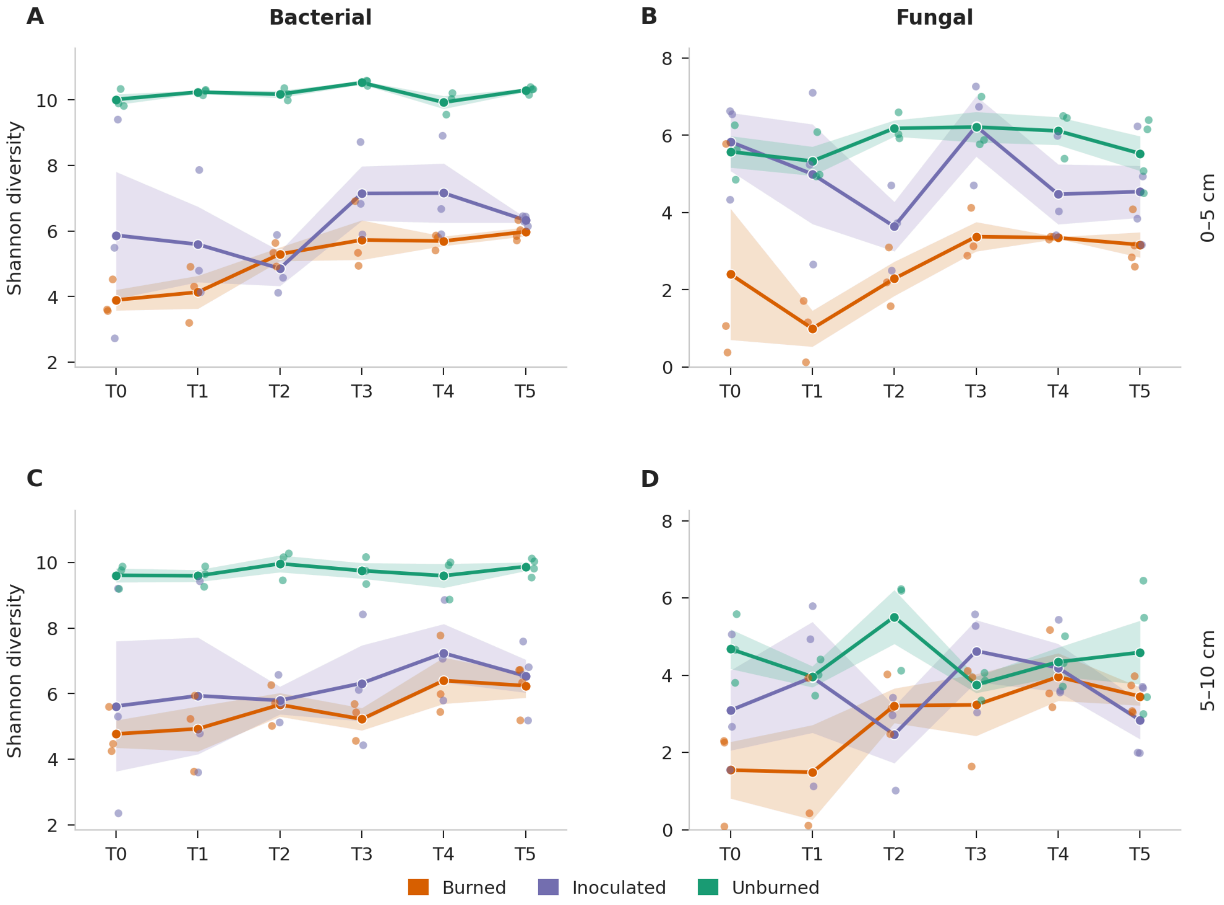


**Figure S7.** Bacterial and fungal alpha diversity across treatments and depths over the 12-month sampling period, estimated from 16S rRNA (A, C) and ITS (B, D) amplicon sequencing at 0–5 cm (A, B) and 5–10 cm (C, D) depth. Lines connect mean Shannon diversity values for each treatment at each time point; shaded bands indicate ±1 standard error; individual points show per-subplot values (n = 3 per treatment at T0–T4, n = 4 at T5). At 0–5 cm, bacterial diversity was substantially reduced in burned and inoculated plots relative to unburned soils at T0 and remained significantly lower throughout the study period (one-way ANOVA with Bonferroni-corrected pairwise t-tests, $p < 0.05$ at all time points). Inoculated plots maintained consistently higher bacterial diversity than burn-only controls, though this difference did not reach statistical significance after correction. Surface fungal diversity showed a similar pattern, with burned plots significantly depleted relative to unburned soils from T1 onward; inoculated plots were intermediate and not significantly different from either burned or unburned soils at most time points. At 5–10 cm, bacterial diversity mirrored the surface pattern, with unburned soils maintaining higher diversity than burned and inoculated plots; significant differences between unburned and burned soils were detected at T2–T5 (Bonferroni-corrected $p < 0.05$), while T0 and T1 did not reach overall significance despite a trend in the same direction. Burned and inoculated plots were not significantly different from one another at any time point. Fungal diversity at 5–10 cm showed no significant treatment differences at any time point after correction, in contrast to the surface.

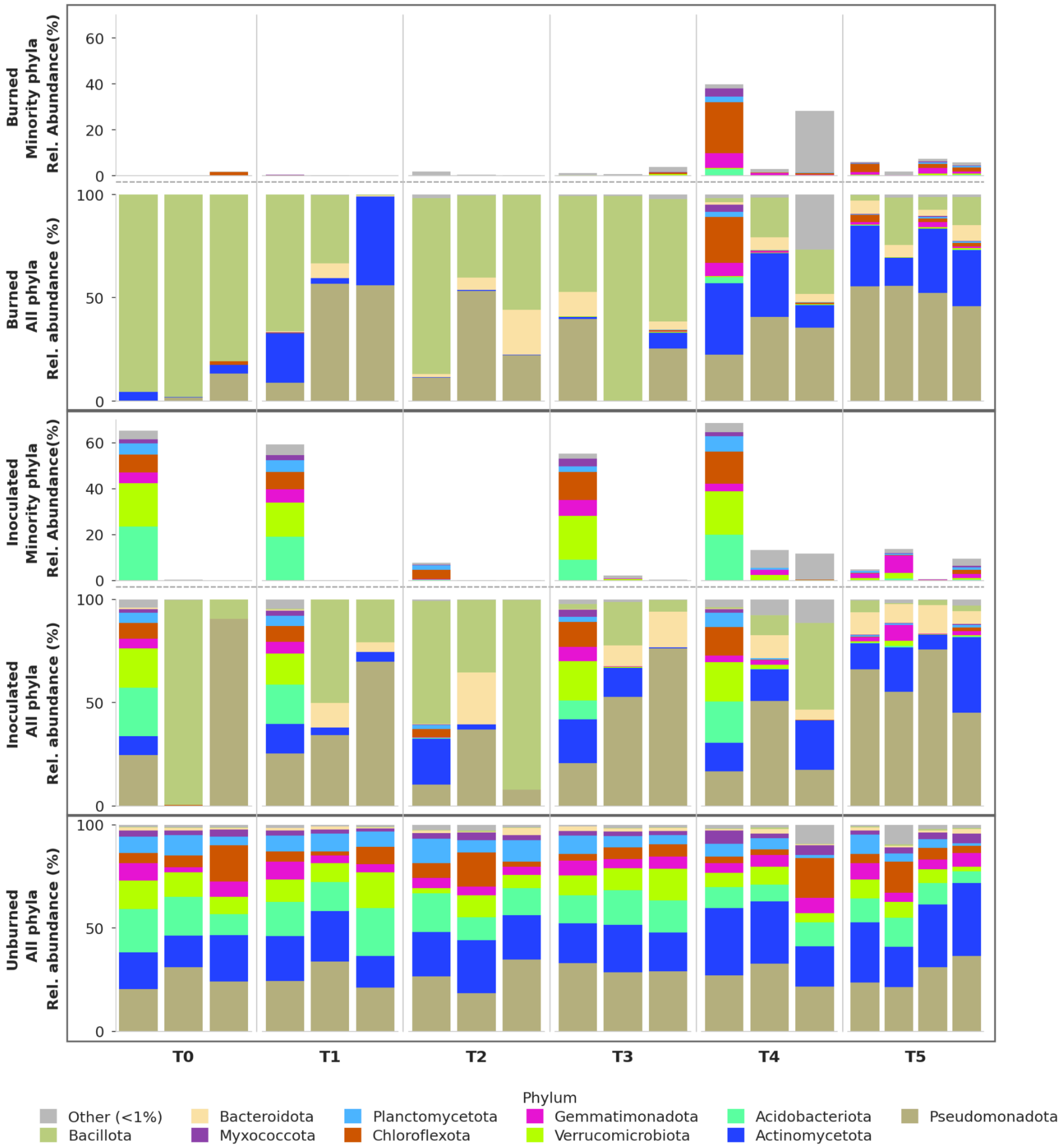


**Figure S8.** Bacterial community composition at the phylum level across treatments and time points (5–10 cm). (A) Stacked bar plots show relative abundance of major bacterial phyla estimated from 16S rRNA amplicon sequencing for each subplot, grouped by condition (unburned, burned, inoculated) and time point (T0–T5). (B) Less dominant phyla, nearly absent from burned plots at T0, begin to return at depth at T4 and T5. Spatially heterogenous recovery is observed in inoculated plots beginning at T0. Unburned soils maintained a compositionally diverse community throughout study.

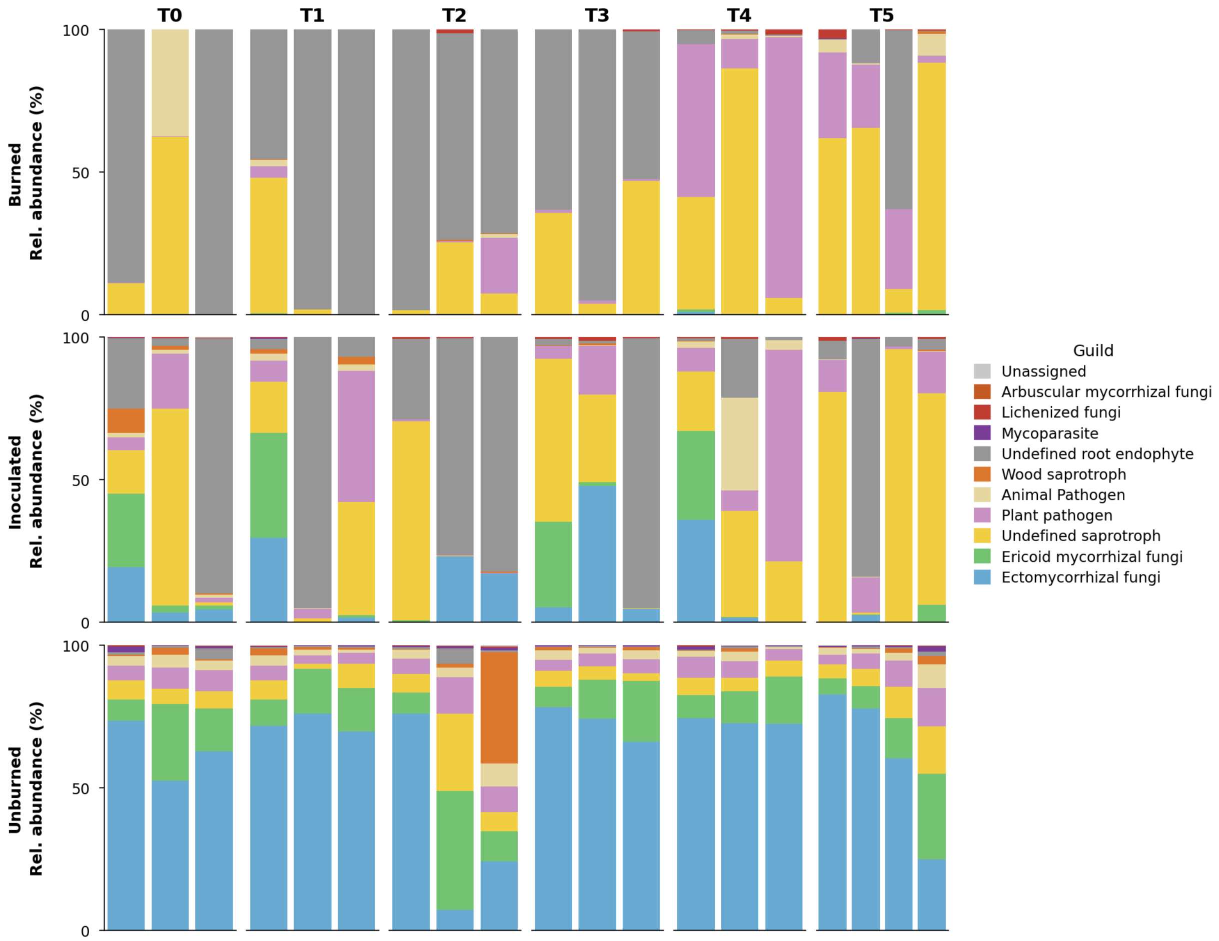


**Figure S9.** Fungal guild composition and ectomycorrhizal community dynamics across treatments at 5–10 cm depth. Stacked bar plots show relative abundance for each subplot replicate (n = 3 per treatment at T0–T4, n = 4 at T5), estimated from ITS amplicon sequencing with FUNGuild annotation (Probable and Highly Probable assignments only). Unburned soils are dominated by ectomycorrhizal (EMF, blue) and ericoid mycorrhizal (ErM, green) fungi throughout the sampling period. Burned plots are dominated by undefined saprotrophs, root endophytes, and plant pathogens. Inoculated plots show restoration of EMF and ErM guilds from T0, with abundance variable across the sampling period.

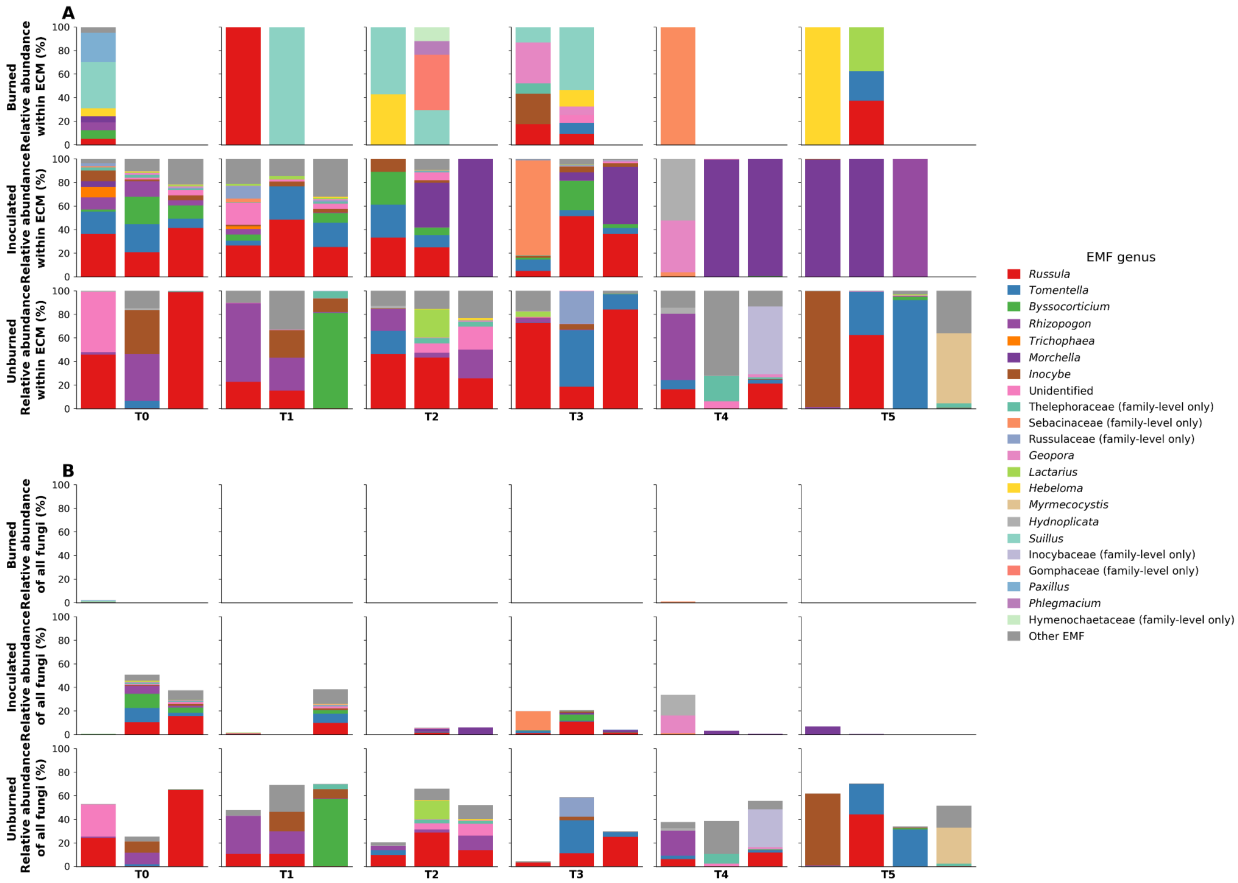


**Figure S10.** Ectomycorrhizal fungal genus composition and temporal dynamics across treatments at 0–5 cm depth. (A) Relative abundance of EMF genera within the total EMF community, expressed as a percentage of EMF ITS reads per sample. (B) Relative abundance of the same EMF genera as a percentage of total ITS reads per sample. In both panels, rows represent treatment (burned, inoculated, unburned) and columns represent sampling time points (T0–T5). Each bar is an individual subplot (n = 3 per treatment at T0–T4, n = 4 at T5). Genera reaching ≥3% mean relative abundance within EMF in at least one treatment are shown individually; the remaining genera are aggregated as "Other EMF." Dominant EMF genera in inoculated and unburned plots include *Russula*, *Tomentella*, *Rhizopogon*, *Inocybe*, *Helvella*, and *Byssocorticium*, mirroring the EMF assemblage of the surrounding intact mixed conifer forest. EMF were near-absent in burned plots throughout the study, as reflected in the very low total EMF signal in panel B.

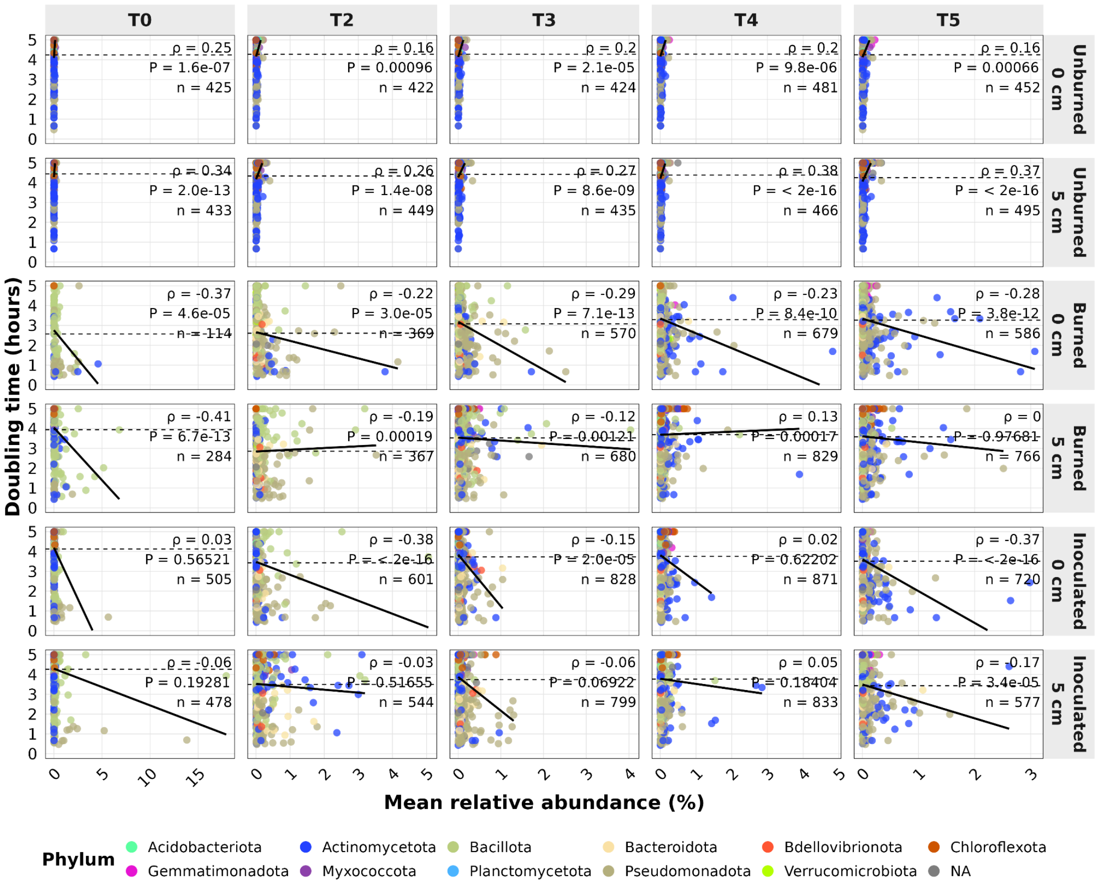


**Figure S11.** Estimated doubling times of organisms across treatments, time points, and soil depths. Scatter plots of mean relative abundance versus estimated doubling time (hours) for all recovered MAGs at 95% ANI, grouped by condition (burned, inoculated, unburned) and time point, at 0–5 cm and 5–10 cm depth. In unburned soils, slow-growing MAGs achieve higher relative abundance (positive slope), consistent with K-selection. This relationship is reversed in burned soils, where fast-growing MAGs dominate (negative slope). These slopes form the basis for the Spearman correlation analysis presented in Figures 3A/B.

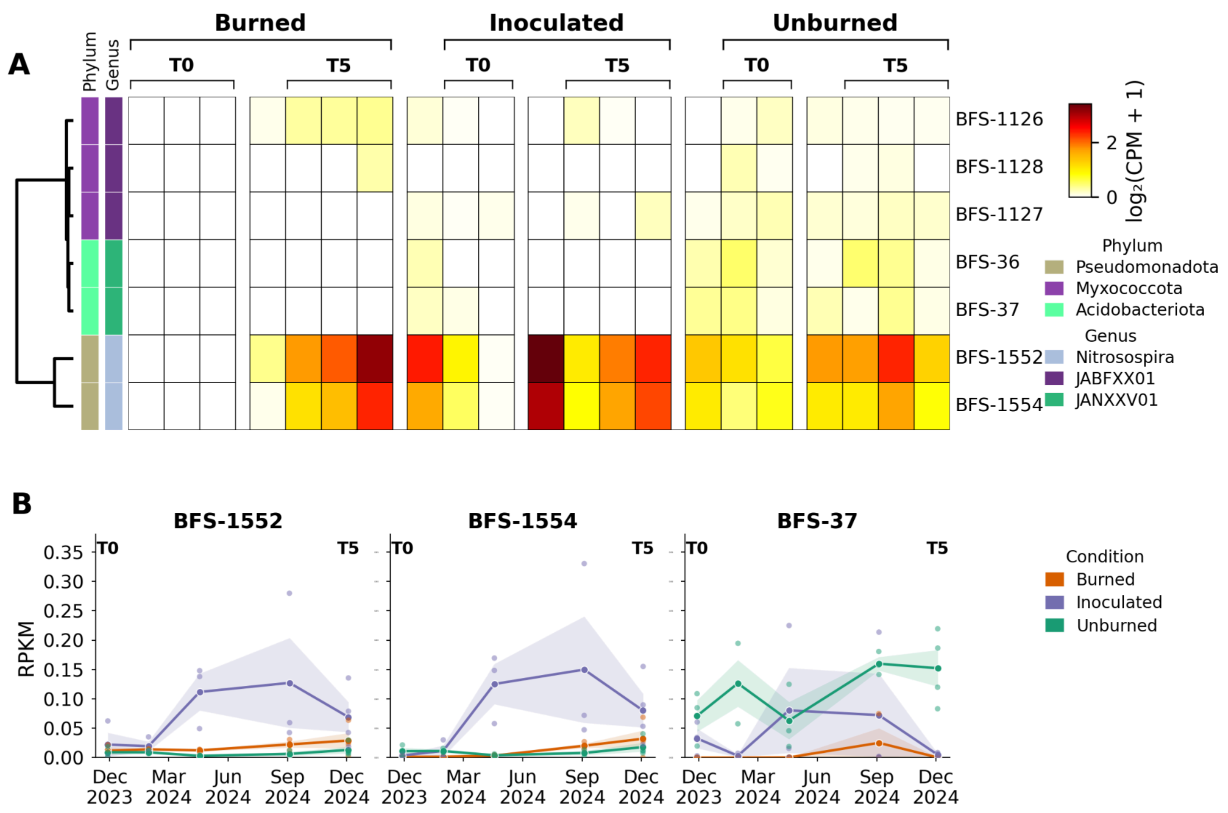


**Figure S12.** MAG-resolved hao transcriptional activity and temporal abundance profiles. (A) Presence and expression of hydroxylamine oxidoreductase (hao) across MAGs for burned, inoculated, and unburned soils at T0 and T5. (B) Relative abundance (RPKM) of select hao-expressing MAGs (BFS-37, BFS- 1552, and BFS-1554) across all sampling time points and conditions.

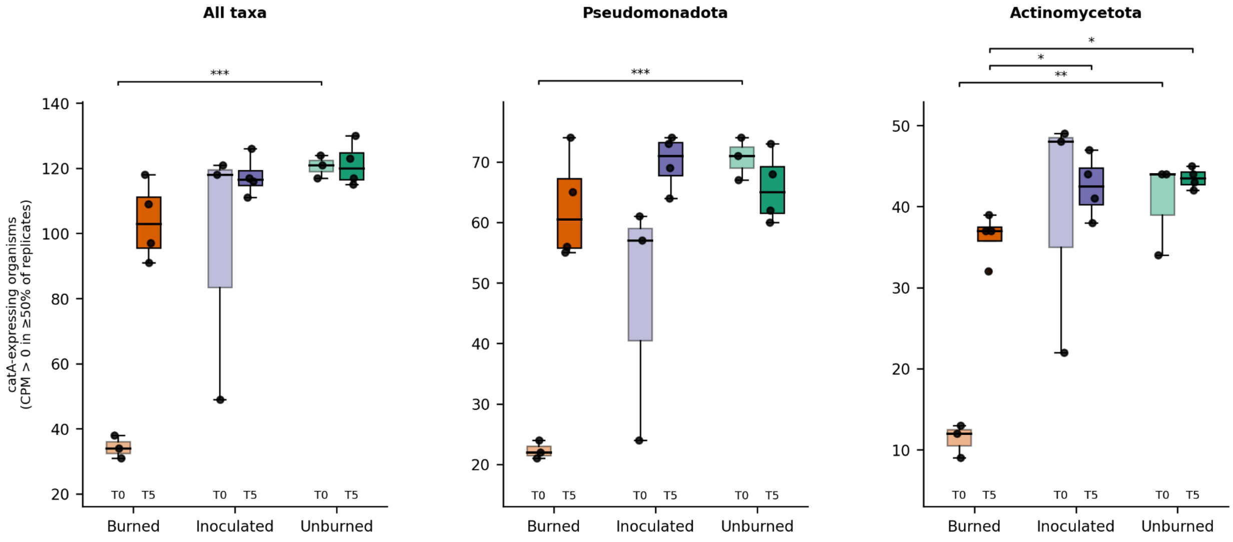


**Figure S13.** Number of organisms with active *catA* transcription (95% ANI MAGs; CPM > 0 in ≥ 50% of replicates) at T0 and T5 across burned, inoculated, and unburned plots, shown for all taxa (A), Pseudomonadota (B), and Actinomycetota (C). Burned soils harbored significantly fewer catA-expressing organisms at T0 compared to unburned soils across all taxa, consistent with fire-induced loss of microbial diversity. Inoculated soils show recovery toward unburned levels from T0, largely driven by Actinomycetota and Pseudomonadota taxa introduced via the inoculant. Pairwise comparisons by Welch's t-test; * $p < 0.05$, ** $p < 0.01$, *** $p < 0.001$.
